# Gene circuit-driven amplified selection enables evolution of fast-growing *Escherichia coli*

**DOI:** 10.64898/2026.08.05.743033

**Authors:** Grayson S. Hamrick, Hye-In Son, Rohan Maddamsetti, Zhengqing Zhou, Kristen Lok, Xiaoli Chen, Aaron Yip, Jing-Mei Qian, Cesar Villalobos, Qianni Ma, Hossein Moghimianavval, Irida Shyti, Ashwini R. Shende, Emma J. Chory, Mary J. Dunlop, Lingchong You

## Abstract

The laboratory *Escherichia coli* K-12 strain has doubled no faster than ∼20 minutes for decades. This plateau could reflect a biophysical limit or simply the way batch culture selects on growth rate. Here we show it can be broken through amplified selection with a Red Queen gene circuit, which takes advantage of growth rate heterogeneity in monoclonal populations to selectively suppress slow-growing cells and creates a tunable mapping from intrinsic growth rate to survival. After 70 days (∼1,000 generations) of amplified selection in MG1655+FHr and subsequent removal of the circuit, a top evolved clone (RQ70) reached a maximum specific growth rate of 2.61 h⁻¹ in shake-flask culture. This corresponds to a doubling time of 15.9 minutes, to our knowledge the shortest reported for *E. coli* K-12, against 18.1 minutes for evolved controls and 20.3 minutes for the ancestor. The gain came at the cost of a ∼3-fold increase in lag time, indicating that the 20-minute plateau is a multi-trait optimum under conventional batch selection rather than an absolute constraint. We argue that synthetic gene circuits can therefore reshape the evolutionary process itself, pushing performance beyond apparent physiological limits.

## Introduction

Growth rate is a fundamental physiological trait that is central to all aspects of microbial physiology, ecology, and evolution.^1–9^ For example, faster-growing cells are often more susceptible to antibiotics that target DNA replication, cell wall synthesis, or protein production, whereas slower-growing or dormant cells can exhibit increased tolerance or persistence.^10–13^ Growth rate is also tied to adaptation to stress factors such as temperature fluctuations, nutrient limitation, and oxidative stress, with survival under adverse conditions often requiring shifts to slower growth.^1,3–5,14–17^ Differences in growth rate drive competitive dynamics among microbial populations by determining how efficiently organisms exploit shared resources, occupy ecological niches, and respond to the presence of competitors or cooperative partners, ultimately shaping community composition and stability.^18–22^

Theoretical studies, grounded in constraints on ribosome biogenesis, macromolecular crowding, and the allocation of cellular resources toward protein synthesis, predict a lower bound on bacterial doubling time of approximately 6 to 9 minutes under idealized conditions.^23–26^ Experimental efforts to approach this limit have identified *Vibrio natriegens* as the fastest-growing free-living bacterium reported to date, achieving doubling times on the order of 10-15 minutes in optimized media, thereby approaching these theoretical predictions.^27–30^

By contrast, *Escherichia coli*, the workhorse of molecular biology and a model for microbial physiology, genetics, and synthetic biology, exhibits substantially slower growth.^31–33^ Under rich, well-aerated conditions, modern bulk-culture studies generally place the doubling time of *E. coli* K-12 near 20 minutes, although isolated values of approximately 17-18 minutes have been reported for other strains or in specific assay configurations.^34,35^ Older literature reports still shorter values, but these results do not provide standardized strain identification, experimental documentation, biological replication, or uncertainty estimates needed for direct comparison with contemporary measurements.^36,37^ This persistent gap between predicted limits, observed maximal growth in other organisms, and the realized growth rate of *E. coli* motivates a central biological question: can the growth rate of *E. coli* be accelerated beyond its currently reported limits, or do fundamental physiological constraints restrict its replication speed?

Addressing this question requires considering how fitness is defined. Selection acts on differences in total reproductive output across the full growth cycle, not on exponential growth rate alone.^38,39^ In serial batch culture, fitness is a composite of multiple cellular traits, including lag time, exponential growth rate, and yield, all of which contribute to total reproductive success across a growth cycle.^40^ Although mutations that increase exponential growth rate do arise, their net selective advantage is often attenuated by trade-offs in other life-history traits and by the fact that exponential growth occupies only part of the growth cycle.^41^

These considerations suggest that the apparent growth-rate plateau in *E. coli* may not reflect an absolute physiological limit, but rather a limitation imposed by the efficiency of selection for exponential growth rate. We hypothesized that the adaptation of growth rate could be accelerated by amplifying the fitness consequences of growth rate variation within a population. To this end, we developed a strategy we term amplified selection – using a synthetic gene circuit to convert small differences in a physiological trait into large differences in survival, greatly strengthening selection on that trait. Here, we implement amplified selection using an engineered “Red Queen” gene circuit that selectively suppresses slow-growing cells within a population, deployed within an experimental design that combines circuit-mediated selection with lineage selection for the fastest-growing lineages. Using amplified selection, we generated an *E. coli* K-12 derivative that doubles every 15.9 minutes during exponential growth, which, to our knowledge, is the shortest doubling time reported for a K-12-derived strain. By engineering both the phenotype-to-fitness mapping and the selection regime, our work demonstrates a generalizable strategy for producing cellular phenotypes that lie beyond established physiological plateaus.

## Results

### An engineered Red Queen (RQ) circuit selectively suppresses slow-growing cells

The circuit (**Figure 1A**) consists of two plasmids: a low-copy (SC101), pCas9 plasmid encoding chloramphenicol (Cm) resistance and an ssrA-tagged endonuclease (Cas9) inducible with anhydrotetracycline (aTc), and a high-copy (pUC) pTarget plasmid carrying both an aTc-activated guide RNA (gRNA) and a constitutive essential survival gene, β-lactamase (AmpR) (see **Supplementary Figure 1** for a detailed plasmid-level schematic). The circuit is turned on by aTc. Through the choice of promoter, ribosome binding site, and Cas9 stability via the addition of an ssrA degron tag, the circuit is tuned such that Cas9 cannot accumulate to a high level in fast-growing cells due to fast growth-mediated dilution.^42^ In contrast, Cas9 will accumulate in slow-growing cells and lead to the cutting of pTarget (**Figure 1B**). Consequently, these cells are selectively suppressed or killed in the presence of carbenicillin (Carb). As a control for the designed circuit function, we also generated a circuit (non-RQ or NRQ) where the target plasmid carries a non-functional gRNA, such that Cas9, even if it accumulates to a high level, cannot cut the target plasmid.

**Figure 1.**
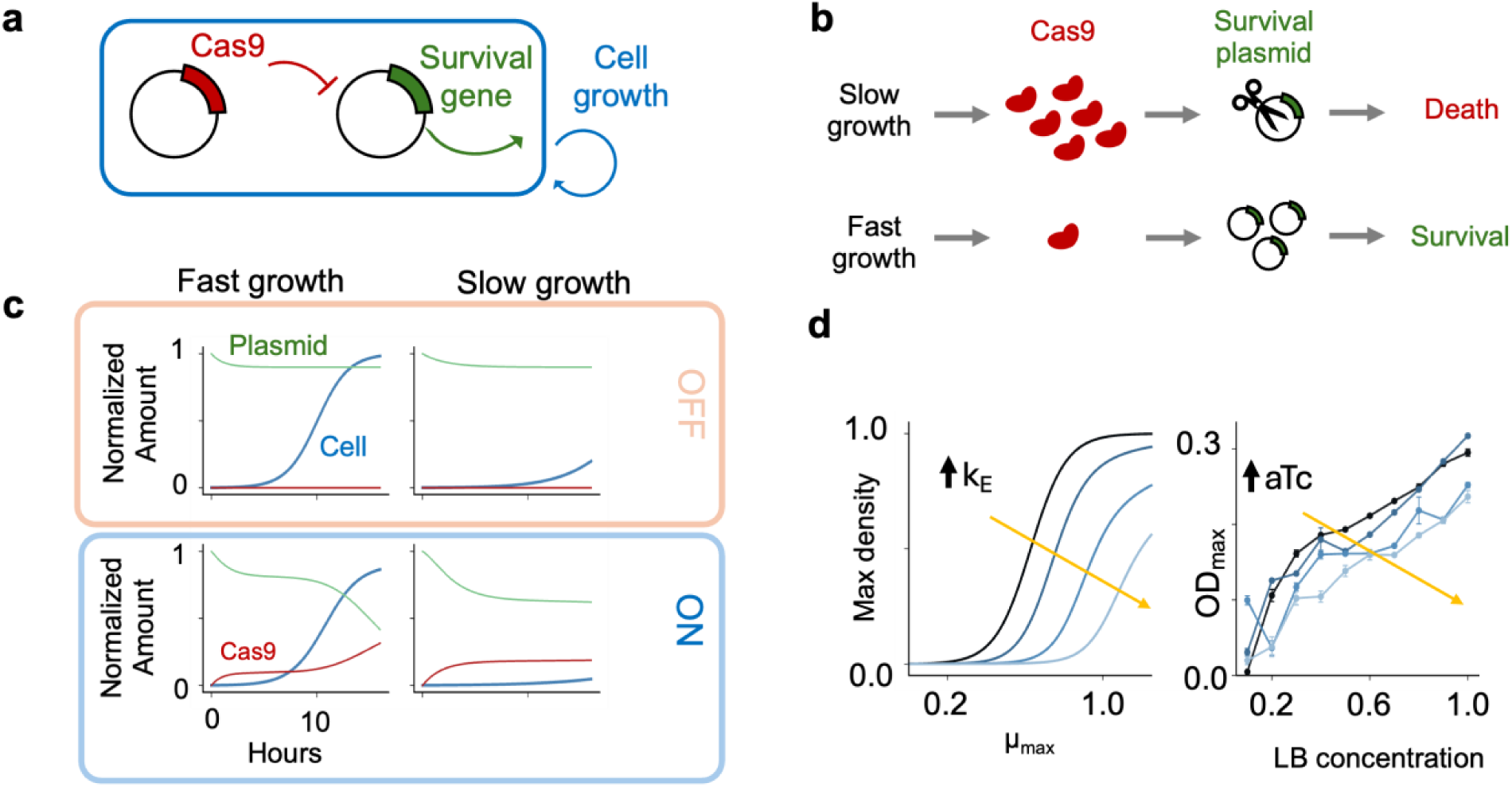
Red Queen (RQ) circuit design. **(a)** The RQ circuit consists of two plasmids: pCas9 and pTarget. pCas9 encodes the endonuclease Cas9, which targets and cleaves a survival plasmid (pTarget). pTarget encodes a beta-lactamase, which allows cell survival in the presence of a beta-lactam antibiotic (e.g. carbenicillin). **(b)** Slow cell growth results in greater accumulation of Cas9, cutting of pTarget, and subsequently cell killing in the presence of carbenicillin. Conversely, rapid growth prevents Cas9 accumulation and subsequent cutting of pTarget, allowing cell survival. **(c)** Model prediction of RQ circuit function. Blue, green, and red curves represent cell population size, pTarget copy number, and intracellular Cas9 concentration, respectively. **(d)** Steady-state cell population across growth rates. Lighter blue curves represent stronger circuit activation (increased *k_E_*). These results are corroborated by experimental evidence, with *μ_max_* varying as a function of LB concentration and induction level varying with aTc concentration.

The circuit design also incorporates a mild negative feedback loop: Cas9-mediated cleavage of pTarget reduces the intracellular abundance of the gRNA cassette, which gradually attenuates further Cas9 activity and prevents runaway plasmid depletion. To balance basal Cas9 cytotoxicity with sufficient dynamic range for circuit modulation, we placed pCas9 on a low-copy SC101 replication origin and pTarget on a high-copy pUC origin. Earlier iterations using a medium-copy ColE1 origin for pTarget produced insufficient β-lactamase to sustain cell growth even with the circuit OFF, resulting in rapid circuit loss during initial testing (**Supplementary Figure 2A**). The final SC101/pUC configuration grew robustly on Cm + Carb plates with and without aTc induction (**Supplementary Figure 2B**), establishing the plasmid backbone pairing used throughout this study.

To gain insight into the circuit dynamics, we developed a kinetic model (**Equations S1-S4**) to account for three key factors: (1) Plasmid copy number-dependent growth inhibition; (2) Cas9-mediated plasmid digestion; and (3) growth-mediated pTarget and Cas9 dilution. In the simulation, the degree of circuit activation is tuned by *κ_E_*, the synthesis rate of Cas9, which reflects aTc-mediated induction in experiments. Consistent with our intuition, the model predicts an amplified dependence of the final cell density of an RQ population on the maximum growth rates (μ_max_) (**Figure 1C**). At a fast growth rate (μ_max_ = 1), the final cell density with circuit fully ON (*κ_E_* = 0.1) is similar to that with circuit OFF (**Figure 1C, bottom left**). At this growth rate, rapid cell division dilutes intracellular Cas9 faster than it can accumulate (red curve), keeping endonuclease concentrations below the threshold required for efficient pTarget cleavage. Consequently, pTarget copy number (green curve) remains high, sustaining β-lactamase expression and permitting robust population growth comparable to the circuit OFF condition. At μ_max_ = 0.5, however, the RQ circuit led to suppression of population growth (**Figure 1C, bottom right**). At this reduced growth rate, growth-mediated dilution is insufficient to clear Cas9, which accumulates to high intracellular concentrations (red curve). The resulting endonuclease activity progressively degrades pTarget (green curve), depleting β-lactamase expression and rendering cells unable to survive carbenicillin selection, causing the population to collapse. The selective suppression is tunable by *k_E_* (**Figure 1D, left**). At increasing *k_E_* values, the population must maintain an increasingly high μ_max_ to achieve a high density at 24 hrs (**Figure 1D, blue curves**). This simulation result is corroborated by experimental evidence that increasing induction of the circuit by aTc decreases yield across concentrations of LB media, a modulator of maximum growth rate (**Figure 1D, right**).

To guide experimental testing, we simulated a two-stage growth process (**Figure 2A**) to assess the circuit’s ability to eliminate cells with insufficient growth rates. In the first stage, cells were cultured until they reached the stationary phase, where the effective growth rate drops to near zero. With the circuit OFF, the model predicts robust growth in both phases (**Figure 2B, left**). To test the predicted function shown in **Figure 2B**, we streaked RQ-carrying *E. coli* Top10F’ cells on LB agar supplemented with 25 µg/mL Cm (selecting for pCas9) and 100 µg/mL Carb (selecting for pTarget), with or without 100 ng/mL aTc. After overnight incubation at 37°C, we inoculated colonies from these plates into LB containing both antibiotics (Cm + Carb) or Cm alone ( **Figure 2B, right**). Cultures derived from RQ colonies with the circuit OFF (no aTc) grew robustly in both media conditions.

**Figure 2.**
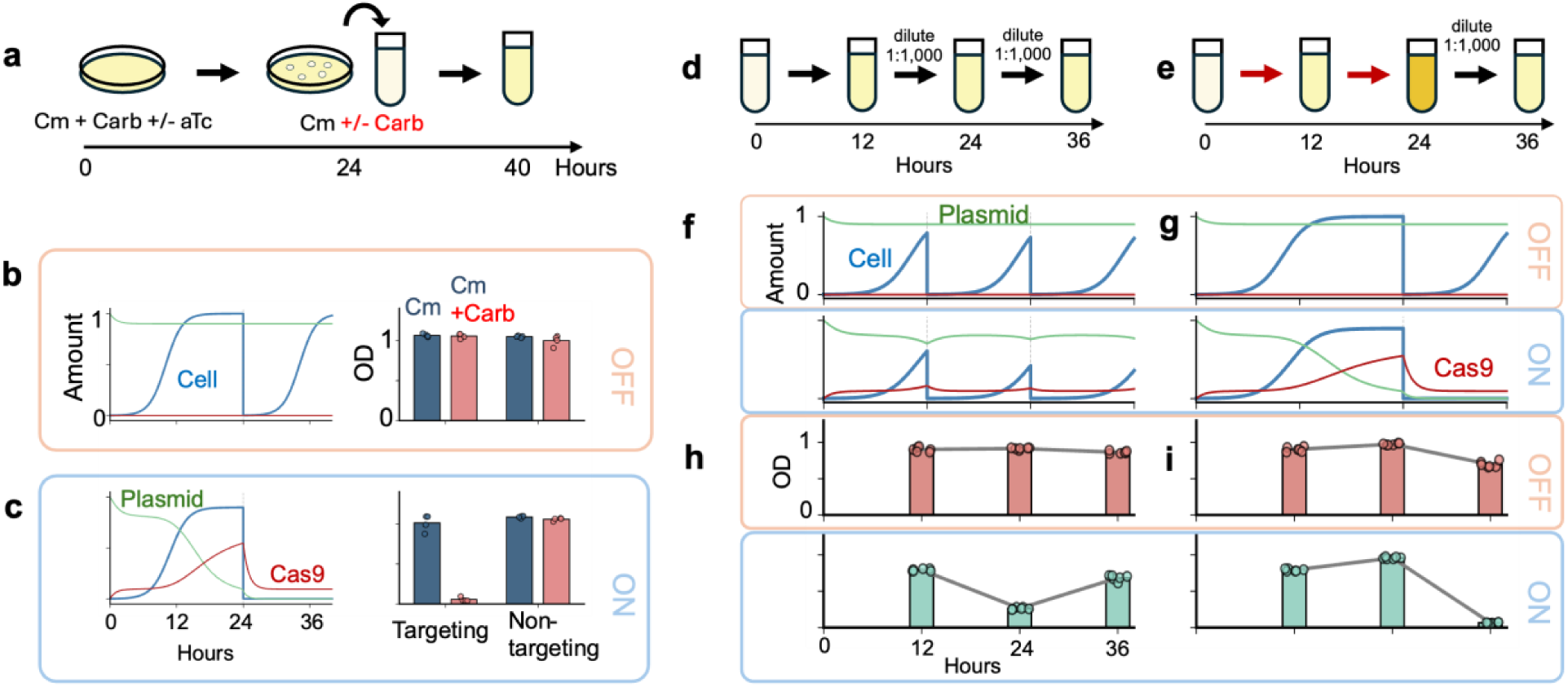
RQ circuit selects against slower growing cells. **(a)** Experimental protocol for agar plate circuit activation. **(b)** Circuit OFF. Modeling (left) of cell population (blue), intracellular pTarget (green) and Cas9 (red) concentrations. Experiment (right). The bar plot shows mean OD600 values from four biological replicates, with individual data point represented by circles. **(c)** Circuit ON. Modeling (left) and experiment (right). The bar plot shows mean OD600 values from four biological replicates, with individual data point represented by circles. **(d)** Experimental protocol for liquid media circuit activation. **(e)** Control experimental protocol. **(f)** Model predictions for cells diluted every 12 hours. **(g)** Model predictions for cells grown for 24 hours before dilution. **(h-i)** Experimental results confirmed the model predictions. Bar plots represent the mean OD600 values from six biological replicates, with individual data points represented by circles. A gray line is included to help visualize OD600 changes over time.

With the circuit ON, the model predicts robust growth during the first phase but population extinction during the second phase (**Figure 2C, left**). This behavior coincides with the accumulation of Cas9 (red) and, correspondingly, cutting of pTarget (green). The simulated population eventually went extinct due to high Cas9 accumulation during the stationary phase before dilution. Cultures derived from RQ colonies exposed to aTc grew in Cm but not in Cm+Carb (**Figure 2C, right**), consistent with predicted circuit function. Cultures derived from non-targeting RQ colonies grew robustly under all conditions (**Figure 2B, C, right**), indicating that Cas9-mediated cutting was critical for the RQ circuit function.

To further evaluate RQ-mediated suppression of slow-growing cells, we simulated and implemented two different culturing processes (**Figure 2D, E**). In one (**Figure 2D**), the culture was diluted every 12 hours to ensure that, on average, the culture maintained a high growth rate between passages. In the other (**Figure 2E**), the culture was not diluted until the third period, such that it would experience a prolonged stationary phase during the first 24 hours. With circuit OFF (*κ_E_* = 0, **Figure 2F, G, top**), the model predicts that the population maintains robust growth (blue) at each stage, which coincides with lack of Cas9 accumulation (red) and plasmid cutting (green).

With circuit ON, however, the two processes led to different outcomes (**Figure 2F, G, bottom**). The cultures maintain robust growth if they are diluted often (every 12 hours) (**Figure 2F, bottom**). In these cultures, Cas9 is maintained at a sufficiently low level (red) due to, on average, sufficiently fast growth of these cultures. Correspondingly, plasmid cutting (green) is negligible. In contrast, if the cultures are not diluted until the 24^th^ hour, they are predicted to crash upon a new passage (**Figure 2G, bottom**). This crash results from the prolonged accumulation of Cas9 (red) and plasmid cutting (green) between 12 and 24 hours (**Figure 2G, bottom**).

Experiments validated these predictions. With circuit OFF, RQ cultures grew throughout following both protocols (**Figure 2H, I, top**). In contrast, with circuit ON (aTc = 100 ng/mL), RQ cultures survived throughout if diluted during each passage (**Figure 2H, bottom**) but crashed if not diluted until the 24^th^ hour (**Figure 2I, bottom**).

### Modeling predicts a biphasic relationship between circuit induction and evolutionary outcome

To explore how amplified selection shapes evolution over serial passages, we simulated 5,000 competing cell lineages, each with a unique growth rate drawn from a narrow initial distribution, all with a shared carrying capacity (**Equations S5–S8**). After each 24-hour passage, a stochastic bottleneck resampled the population proportional to final abundance, and daughter lineages acquired small heritable growth-rate variation (*σ* = 0.01*μ_max_*_,*parent*_). Populations whose surviving fraction fell below 0.01% were considered extinct.

Without the circuit (adaptive evolution), natural competition increased the population mean growth rate by ∼0.29 h⁻¹ over 25 passages (**Figure 3A**). With amplified selection at *κ_E_* = 1.0, the gain was ∼0.44 h⁻¹, a 1.5-fold increase relative to adaptive evolution (**Figure 3A**). As the mean population growth rate increased, the surviving population fraction with each passage increased as well (**Supplementary Figure 3A**).

**Figure 3.**
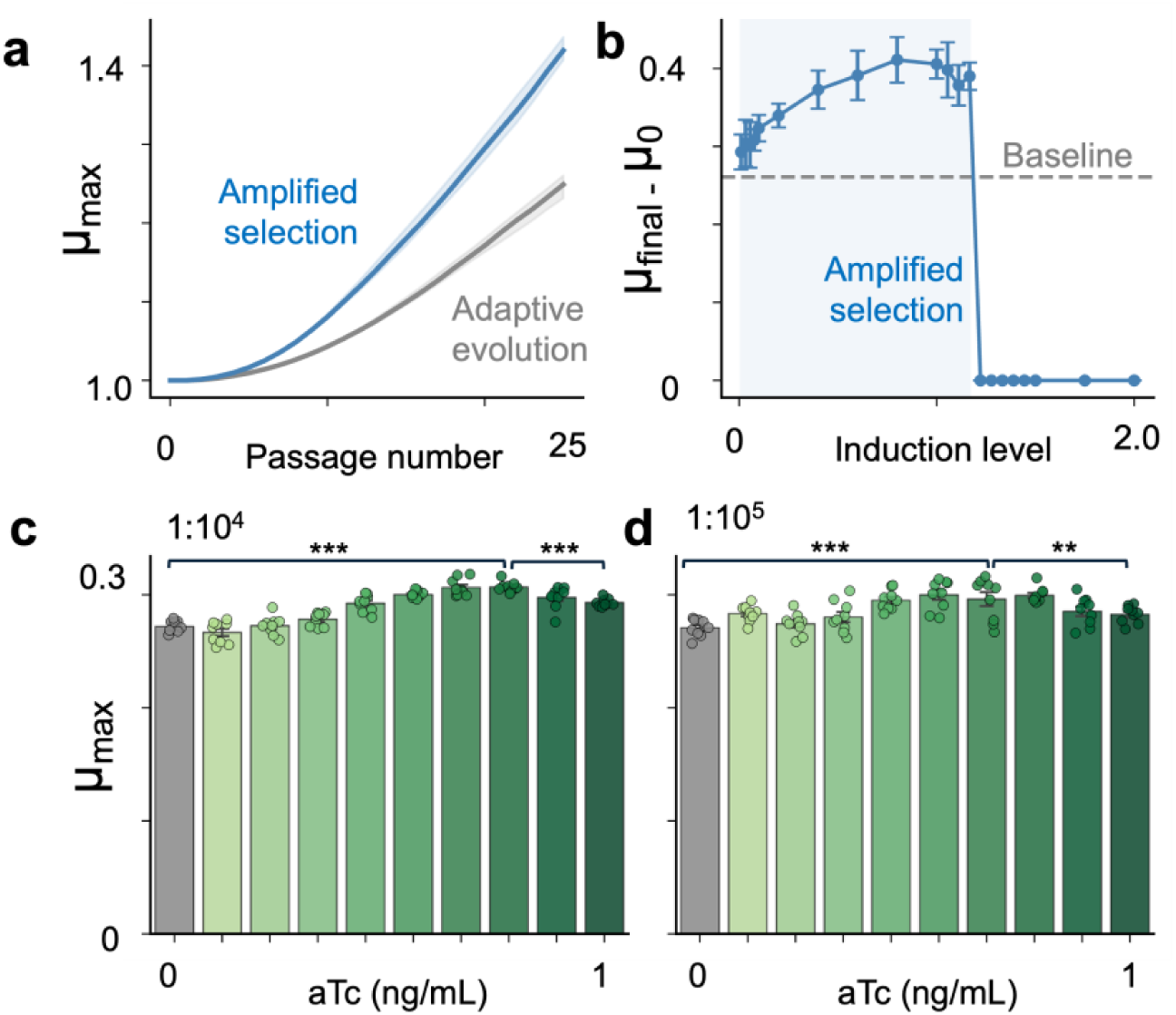
Amplified selection accelerates growth-rate improvement but exhibits a biphasic dependence on induction level. **(a)** Mean population growth rate over 25 serial passages for cells with (blue) and without (gray) amplified selection at *k_E_* = 1.0. Shading indicates interquartile range across replicates. **(b)** Selection gain (final mean growth rate minus initial) as a function of *k_E_*. Dashed gray line indicates the baseline improvement without amplified selection. The shaded region marks the *k_E_* range where amplified selection exceeds baseline. Error bars indicate ± 1 s.d. across replicates. **(c)** aTc-inducible growth-rate titration in MG1655+FHr RQ cells at Day 3 with 1:10,000 daily dilution. ***: p < 0.001. **(d)** aTc-inducible growth-rate titration in MG1655+FHr RQ cells at Day 3 with 1:100,000 daily dilution. **: p < 0.01.

Sweeping *κ_E_* across two orders of magnitude (0.01–2.0) revealed a biphasic relationship between induction level and selection performance (**Figure 3B**). At low *κ_E_*, the circuit provided marginal benefit over adaptive evolution. Selection gain increased with *κ_E_* through intermediate values, peaking near *κ_E_* = 1.1. Above *κ_E_* = 1.2, all replicates went extinct within the first passage. Mean per-passage survival declined monotonically with *κ_E_*, from ∼98% at *κ_E_* = 0.01 to ∼5% at the optimum *κ_E_* = 1.1 (**Supplementary Figure 3B**).

### Intermediate induction of the RQ circuit maximizes growth rate during short-term serial passaging

To ask how the level of circuit induction shapes the growth phenotype that emerges under amplified selection, we serially passaged MG1655+FHr carrying the RQ circuit for two days across a range of aTc concentrations (0–1 ng/mL) at two daily dilution factors. On Day 3, we determined the maximum specific growth rates from plate-reader growth curves (**Figure 3C and Figure 3D**) by fitting each to a logistic model incorporating a lag phase (**Equations S9**).

Induction increased the acquired growth rate relative to the uninduced control across both dilutions, but the dependence on aTc was non-monotonic: growth rate rose to a plateau between roughly 0.1 and 0.6 ng/mL and declined at saturating induction. At the 1:10,000 daily dilution, the plateau maximum exceeded both the uninduced control (Δμ = 0.035 h⁻¹) and the saturating dose (Δμ = 0.014 h⁻¹; both Welch p < 0.0001). The same pattern held at 1:100,000, where the plateau maximum again exceeded both the uninduced control (Δμ = 0.029 h⁻¹, p < 0.0001) and the saturating dose (Δμ = 0.017 h⁻¹, p < 0.005). Concentrations within the plateau were mutually indistinguishable (Tukey HSD p > 0.13), indicating a broad optimum rather than a sharply tuned peak.

This non-monotonic dose–response mirrors the biphasic dependence on induction strength (*κ_E_*) predicted by our amplified selection model (**Figure 3B**), consistent with induction levels beyond an optimum imposing a cost that offsets the selective benefit.

### Amplified selection accelerates E. coli Top10F’ growth rate over 100 days

To test whether RQ-mediated amplified selection could drive measurable improvements in growth rate over extended serial passaging, and to establish that circuit function generalizes across laboratory strain backgrounds, we first conducted a proof-of-concept long-term evolution experiment in *E. coli* Top10F’ spanning 100 days (**Supplementary Figure 4A**). Four lineages of RQ cells carrying a targeting spacer (sp), two lineages carrying a non-targeting spacer (NT), and three *E. coli* Top10F’ lineages without the circuit were passaged daily in parallel. For the first 43 days, cultures were diluted 10,000-fold into LB supplemented with Cm, Carb, and one of four aTc concentrations (0, 0.1, 1, or 10 ng/mL) spanning three orders of magnitude. On Day 43, the two fastest-growing targeting lineages (both evolved in 0.1 ng/mL aTc) and one non-targeting lineage were selected for continued evolution under more stringent selection — a 100,000-fold daily dilution and a 10-fold higher aTc concentration (1 ng/mL) — for an additional 57 days.

By Day 100, the two selected targeting RQ lineages exhibited 1.95- and 1.86-fold increases in maximum growth rate relative to the ancestor, reaching 0.57 and 0.50 h⁻¹ respectively (**Supplementary Figure 4B, left and middle; Supplementary Figure 5**). The non-targeting control lineage achieved a 1.16-fold increase (0.41 h⁻¹; **Supplementary Figure 4B, right; Supplementary Figure 6**), consistent with adaptation to the plasmid-bearing, antibiotic-containing growth environment in the absence of circuit-mediated selection. The three naïve lineages, passaged in antibiotic-free LB, increased 1.10- to 1.27-fold over the same period (**Supplementary Figure 7**). Evolved targeting RQ lineages thus outpaced both their non-targeting counterparts and the antibiotic-free controls. These growth-rate gains are consistent with a contribution from RQ-mediated amplified selection beyond generic laboratory adaptation.

These results established that amplified selection is robust to extended serial passaging and transfers across *E. coli* strain backgrounds. They also motivated two design refinements for our subsequent effort: (i) that 0.1–1 ng/mL aTc falls within the productive induction window predicted by the biphasic modeling above, and (ii) that circuit-mediated within-lineage selection alone, while effective, leaves room for additional gain if combined with active experimenter-directed between-lineage selection. We therefore developed a more aggressive multi-level selection pipeline in *E. coli* MG1655+FHr, described below.

### *Amplified selection produced an* E. coli *K-12 derivative with a minimum doubling time of 15.9 minutes*

To determine whether RQ-mediated selection could be harnessed to produce an *E. coli* lineage with a substantially faster exponential growth rate, we designed a selection pipeline spanning 70 days of serial passaging. The pipeline combined two levels of selection: circuit-mediated growth-rate discrimination within lineages and experimenter-directed propagation of the fastest-growing lineages (**Figure 4A, inset**). Populations of MG1655+FHr cells without the circuit were passaged in LB medium, while MG1655+FHr cells carrying the RQ circuit evolved under gradually increasing induction by aTc in LB containing chloramphenicol (Cm) and carbenicillin (Carb). We adopted a protocol to better preserve the growth-rate–dependent fitness transformation imposed by the RQ circuit. Overnight cultures were first diluted 1,000-fold and precultured in LB containing Cm and Carb without aTc for 2 hours to bring them to exponential phase and were then diluted 100-fold (for the first 40 days of passaging) or 1,000-fold (for the final 30 days of passaging) into LB containing Cm, Carb, and aTc. Inducing the circuit during exponential growth ensured that survival depended on instantaneous growth rate, thereby maintaining the nonlinear mapping between intrinsic growth and effective fitness. Induction levels were tuned to avoid complete circuit activation. Strong induction (>10 ng/mL aTc) caused severe growth inhibition and frequent circuit disruption, selecting for escape mutants and risking population extinction. Instead, moderate induction (0.1–1 ng/mL aTc) imposed a soft but consequential growth-rate threshold that amplified fitness differences while preserving viability. aTc concentrations were gradually increased over time to maintain selective pressure as populations adapted.

**Figure 4.**
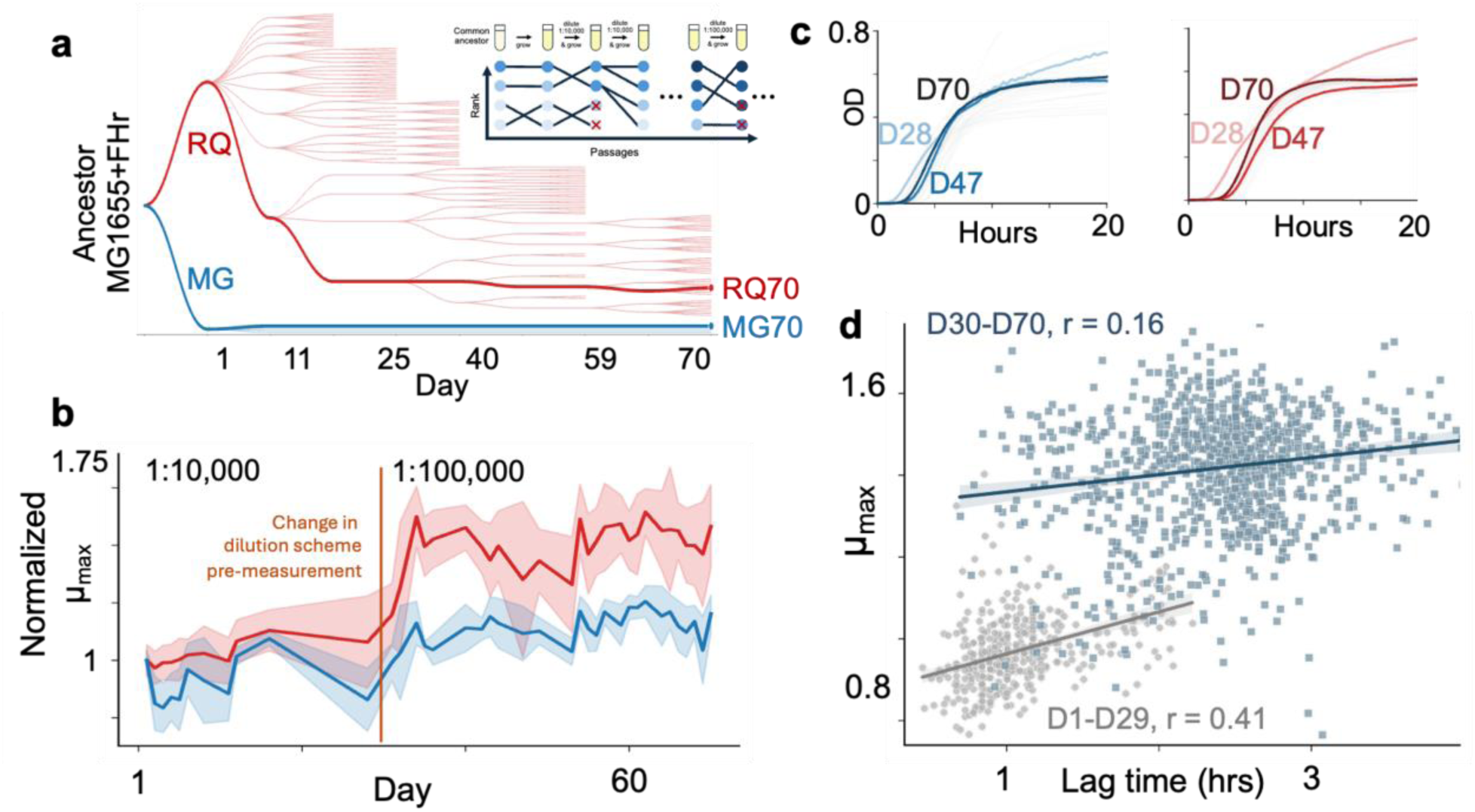
Amplified selection accelerates growth-rate evolution. **(a)** Long-term evolution of MG1655+FHr with or without the RQ circuit. The cladogram shows ancestor MG1655+FHr and its descendants from RQ-driven evolution (red) or adaptive evolution (blue). **(b)** Growth-rate trajectories over the evolution, normalized to the growth rate at Day 1, for RQ (red) and MG (blue) cells. **(c)** Representative growth curves from the longitudinal data collection for the MG70 lineage (left, blue) and the RQ70 lineage (right, red). **(d)** Growth rate and lag are moderately correlated in the first dilution scheme (r = 0.41, n = 447) and very weakly correlated in the second dilution scheme (r = 0.16, n = 1,014).

Growth trajectories were monitored throughout the experiment. Approximately every 14 days, slower-growing lineages were discarded while the fastest-growing populations were expanded into multiple wells, progressively enriching the experiment for high-performing lineages (**Figure 4A**). Under this amplified selection regime, the growth rate *μ_max_*, as determined by fitting a logistic-with-lag growth model to growth curves, increased more quickly compared to the control group undergoing adaptive evolution without the circuit (**Figure 4B**).

By Day 70, the cells had experienced ∼1,000 generations. Longitudinal growth curve measurements were taken approximately every other day. Four control populations without the circuit exhibited gradual improvement, with a mean growth-rate increase of ∼1.2-fold from Day 1 to Day 70 (from 1.28 h⁻¹ to 1.54 h⁻¹) (**Figure 4B**). In contrast, the RQ lineages evolved faster, achieving up to ∼1.6-fold increases in growth rate (from 0.94 h⁻¹ to 1.50 h⁻¹) under moderate induction (0.1–1 ng/mL aTc), while carrying the circuit and under selection by Cm and Carb. This is evident from the qualitative change in the growth curves in **Figure 4C**. Growth rate and lag are moderately correlated in the first dilution scheme, when cells were diluted 10,000-fold prior to measurement (r = 0.41, n = 447) and very weakly correlated in the second dilution scheme, where cells were diluted 100,000 prior to measurement (r = 0.16, n = 1,014) (**Figure 4D**).

To determine whether the growth-rate gains produced by the selection pipeline reflected stable improvements in cellular growth capacity, we selected a clone from the fastest-growing RQ lineage for detailed physiological characterization. After curing of the RQ circuit, confirmed by loss of antibiotic resistance, this clone — designated RQ70 — was evaluated alongside evolved controls (MG70) and the ancestral strain (MG) in shake-flask cultures. We used plating and colony counting to observe growth curves and evaluate growth rates in 20 mL cultures in baffled shake flasks (**Figure 5A**). All measurements were performed in LB medium at 37 °C without antibiotics or inducers. RQ70 exhibited a maximum specific growth rate of 2.61 ± 0.05 h⁻¹, corresponding to a minimum doubling time of 15.9 minutes across 12 biological replicates (**Figure 5B**). This compares to 2.30 ± 0.08 h⁻¹ (18.1 minutes) for MG70 cells (3 biological replicates) and 2.04 ± 0.05 h⁻¹ (20.3 minutes) for the ancestral MG strain (3 biological replicates), as calculated by CFU counting (ANOVA p < 0.0002, with Tukey HSD p < 0.02 between RQ70 and MG70 and p < 0.0002 between RQ70 and ancestral MG). RQ70 therefore crossed the widely reproduced ∼20-minute K-12 benchmark by a significant margin: its doubling time was approximately 22% shorter than that of the ancestor and 12% shorter than that of the evolved control The growth-rate-and-lag trade-off was evident under this condition as well: RQ70 has a lag time (3.03 hours) roughly double that of MG70 (1.42 hours) and almost three times that of the ancestral MG strain (1.12 hours) (ANOVA p < 0.0002, with Tukey HSD p < 0.003 between RQ70 and MG70 and p < 0.0006 between RQ70 and ancestral MG) (**Supplementary Figure 8**).

**Figure 5.**
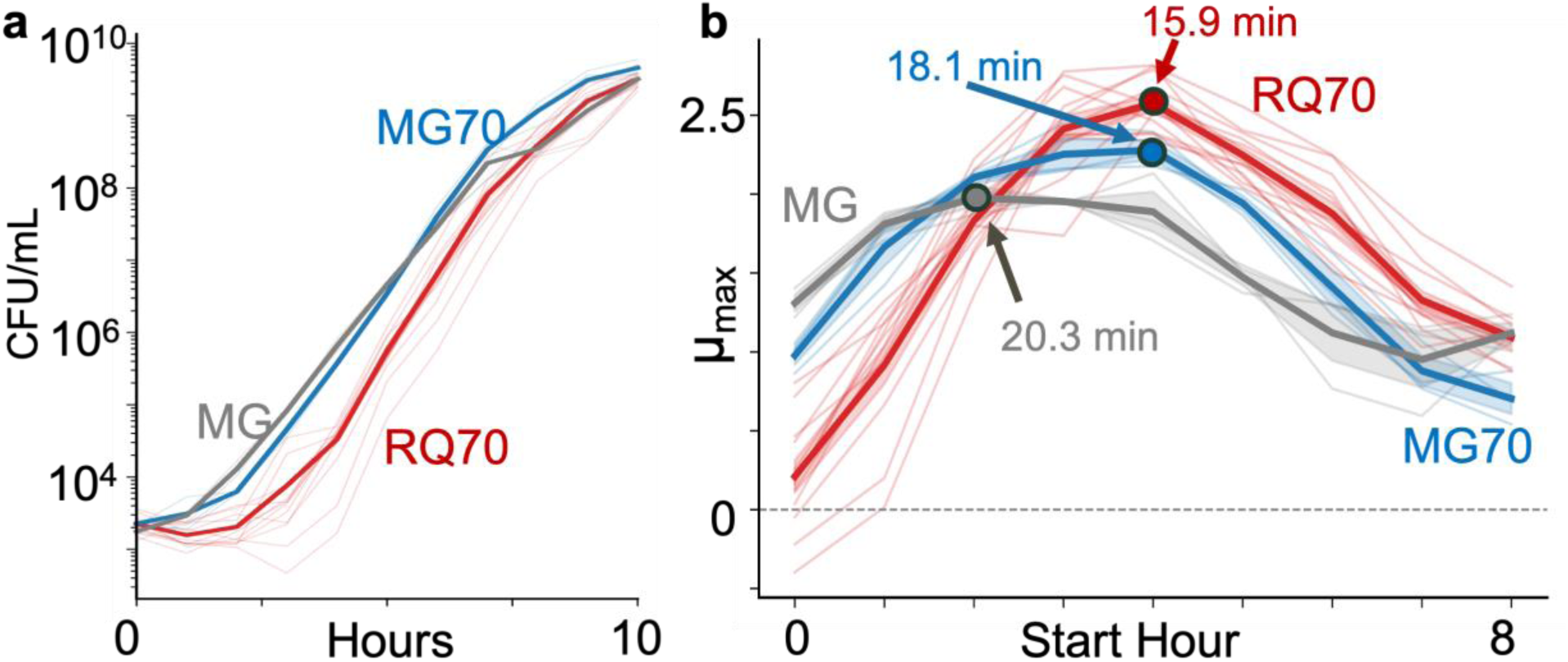
RQ70 achieves a minimum doubling time of 15.9 minutes. **(a)** Shake flask growth curves for RQ70, MG70, and ancestral MG. **(b)** Maximum 3-hour rolling growth rate. Cured RQ70 reached 2.61 ± 0.05 h⁻¹, corresponding to a minimum doubling time of 15.9 minutes, compared with 18.1 minutes for the Day 70 evolved wild-type (MG70) cells and 20.3 minutes for the ancestral MG1655+FHr cells.

### RQ70 experienced significant trade-offs in its phenotypic traits

To examine how RQ70’s growth-rate advantage manifests under continuous culture conditions, we compared RQ70, MG70, and MG cells in turbidostat culture using a microplate-based system across four OD setpoints (0.6, 0.7, 0.8, 0.9; n = 8 biological replicates per condition). Under steady-state exponential growth (t = 5–10 h), RQ70 and the MG70 control achieved indistinguishable growth rates at all setpoints tested (e.g., 0.79 ± 0.02 and 0.79 ± 0.02 h⁻¹ at OD setpoint 0.8, respectively), while both evolved strains exceeded the ancestor (0.60 ± 0.01 h⁻¹; **Supplementary Figure 9**). Growth rate declined with increasing OD setpoint for all strains, consistent with expected turbidostat dynamics. The convergence of RQ70 and MG70 in this context may reflect a shared environmental ceiling imposed by the static microplate format, where oxygen transfer and mixing are limited relative to baffled shake flasks. Regardless of mechanism, both evolved strains maintained a substantial growth-rate advantage over the ancestor (∼1.3-fold at all setpoints), confirming that the selection pipeline produced durable improvements in exponential growth capacity that persist outside of batch culture conditions.

We also measured growth of RQ70, MG70, and ancestral MG cells with single-cell resolution in both a microfluidic mother machine (**Supplementary Figure 10**) and under agarose pads. Under these conditions, RQ70 exhibited the slowest median single-cell growth rate of the three strains (**Supplementary Figures 11-12**).

We further compared RQ70, MG70, and MG in M9 minimal medium with 0.4% glucose (**Supplementary Figure 13**). Under these nutrient-limited conditions the absolute growth rates of all strains were substantially lower than in LB, but the rank order was consistent: RQ70 grew fastest (μ = 0.44 h⁻¹; doubling time ≈ 94 min), followed by MG70 (μ = 0.29 h⁻¹; ≈ 145 min) and the ancestor (μ = 0.21 h⁻¹; ≈ 198 min). RQ70 also established fastest (time to OD 0.05 ≈ 3.0 h, versus 3.2 h for MG70 and 5.1 h for the ancestor) and reached the highest final yield (max OD ≈ 0.59, versus 0.50 and 0.31). The growth traits for RQ70, MG70, and ancestral MG are summarized in Figure 6A.

**Figure 6.**
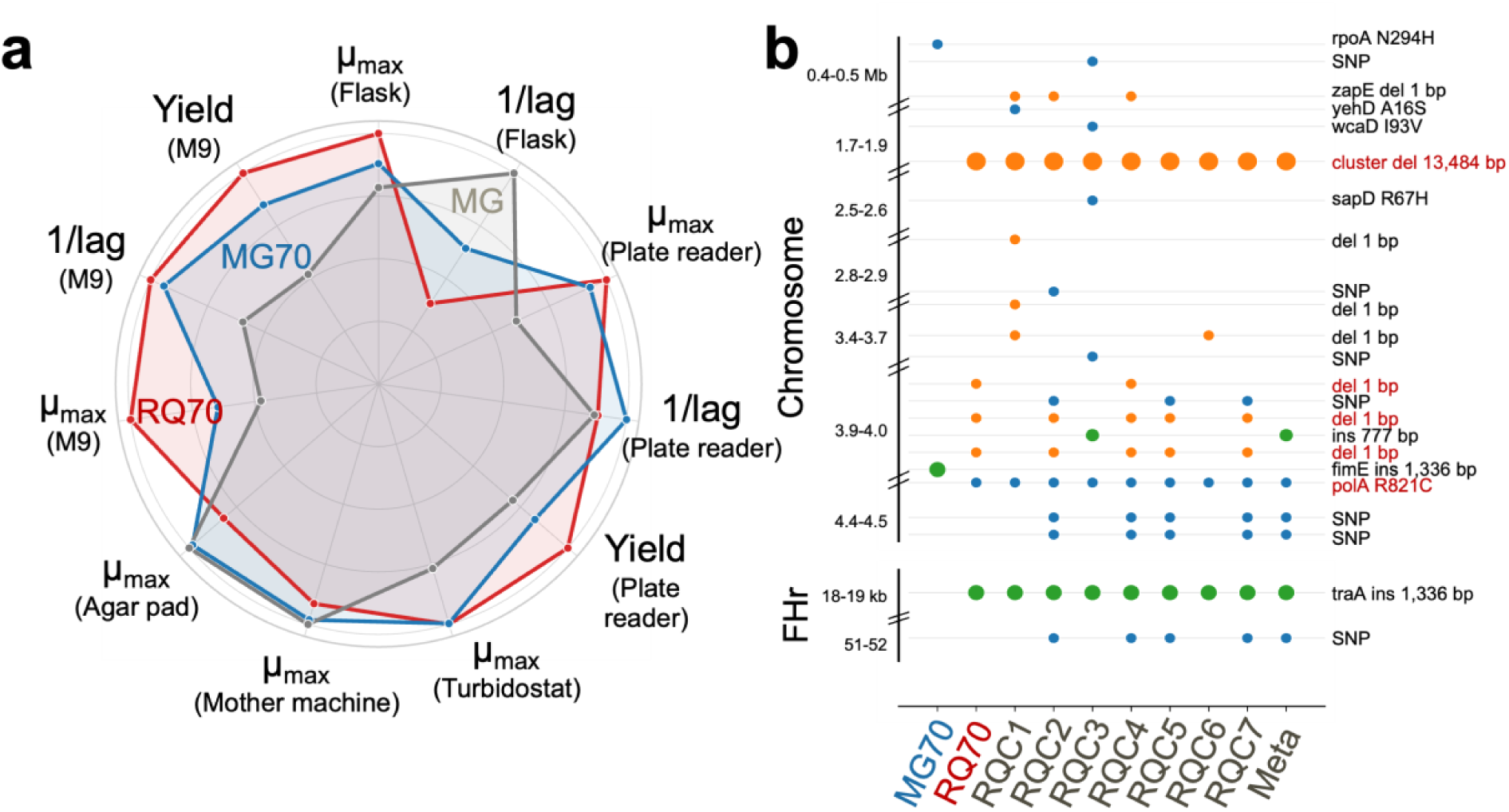
Phenotypic and genomic characteristics of RQ70 and MG70. **(a)** RQ70 phenotype trade-offs relative to MG70 and the ancestor, with each axis normalized to the strain with the maximum value for that attribute. **(b)** Genomic divergences between MG70, RQ70, additional RQ clones within the RQ70 metapopulation, and the ancestral MG background. Blue dots represent SNPs, orange deletions, and green insertions.

Carriage of a kanamycin-resistant plasmid under selection reduced growth in a strain- and copy-number-dependent manner (**Supplementary Figure 14**). Relative to matched M9-only wells, the ancestor and MG70 lost substantial integrated growth — the ancestor retained only 70% (p15A) and 66% (pADEPT) of its AUC and lost 22% and 41% of its yield. MG70 retained ≈ 80% of AUC, whereas RQ70 was essentially unaffected (94–97% of AUC retained; yield ratios ≈ 1.0). The cost was larger for the high-copy pADEPT than for the medium-copy p15A, consistent with a copy-number-dependent burden, and was accompanied by a modest increase in lag (≈ 0.5–1 h) across strains.

The accelerated RQ70 lineage was more sensitive to gentamicin, kanamycin, and chloramphenicol (**Supplementary Figures 15-17**). At intermediate-to-high antibiotic concentrations, the RQ strain exhibited a prolonged lag followed by delayed exponential escape, while at the highest concentrations tested its growth was largely suppressed within the assay window. In contrast, the MG70 strain was more resistant to all three antibiotics than the ancestral MG strain.

### Genomic characterization of RQ70 reveals chromosomal mutations

Whole-genome sequencing of RQ70 revealed several chromosomal mutations and one mutation to the FHr plasmid relative to the ancestral MG1655+FHr reference (**Figure 6B**). The chromosomal mutations to coding regions consisted of a single nucleotide polymorphism (SNP) in the gene for DNA polymerase I (polA; R821C) and a 13,484-base deletion of the flagellar and chemotactic systems spanning 16 genes. The plasmid mutation was a 1,336-base insertion of a transposon disrupting the type IV conjugative transfer system pilin TraA. In contrast, MG70 exhibited a single SNP in the DNA-directed RNA polymerase subunit alpha *rpoA* gene (N294H). Additional clones sequenced from the RQ70-bearing well indicate that the RQ70 mutations were fixed in the population, while additional idiosyncratic insertions, deletions, and SNPs were present elsewhere in the population (**Figure 6B**).

## Discussion

Synthetic gene circuits are generally used to program new cellular behaviors. Here, we instead use a synthetic gene circuit to interrogate the limits of physiological growth in *E. coli*. Rather than accelerate *E. coli* growth directly, the RQ circuit selectively suppresses slower-growing cells and thereby changes the mapping from growth phenotype to survival. Biosensor-coupled selections and phage-assisted continuous evolution similarly link an engineered output to fitness.^43,44^ RQ extends this logic to a native physiological trait and uses the resulting selection to probe the accessibility of faster growth.

In MG1655+FHr, combining circuit-mediated selection within lineages with repeated propagation of the fastest-growing lineages yielded RQ70. After circuit curing, RQ70 reached a maximum specific growth rate of 2.61 h⁻¹, corresponding to a minimum doubling time of 15.9 minutes, compared with 18.1 minutes for MG70 and 20.3 minutes for the ancestor (**Figure 5B**). To our knowledge, this is the shortest doubling time reported for an *E. coli* K-12 strain.

Why did amplified selection reach this state when conventional adaptive laboratory evolution has not? One possibility is that RQ simply accelerates the rise of fast-growing variants that would also be favored during conventional evolution; given sufficient time, a conventional experiment might therefore reach a comparable strain. Alternatively, RQ may change which mutational steps are beneficial. Without the circuit, individual mutations that ultimately contribute to the RQ70 phenotype may provide little or no benefit on their own and therefore fail to establish. In the circuit-bearing state, however, the physiological state created by RQ expression—not necessarily through its intended function alone—could make one such mutation beneficial. Once established, this mutation could create a genetic background in which additional mutations become beneficial. RQ could thereby open an adaptive path to RQ70 that conventional selection is unlikely to traverse. The two possibilities are not mutually exclusive.

Past studies support the possibility that a circuit-defined physiological state could reshape the adaptive landscape. Synthetic circuits can perturb host physiology and generate feedback between circuit activity and cell growth.^45,46^ Environmental and genetic context can, in turn, change the fitness effects of mutations and the mutational paths that are selectively accessible.^47–49^ The evolution of aerobic citrate utilization in the *E. coli* long-term evolution experiment provides a particularly relevant example: the fitness effect of activating *citT* ranged from beneficial to highly deleterious across evolved genetic backgrounds, and a potentiating *gltA* mutation helped restore a background in which this first step could persist until additional mutations refined the phenotype.^50–52^

Two observations are consistent with the possibility that RQ altered evolutionary accessibility. First, RQ70 was not uniformly faster across environments. It outperformed the controls in LB shake-flask and M9-glucose batch cultures, but was indistinguishable from MG70 in turbidostat culture, had the lowest median single-cell growth rate in the mother machine, and grew more slowly than MG70 and the ancestral MG on agar pads (**Figure 5B; Supplementary Figures 9 and 11-12**). These assays differ in nutrient renewal, population density, spatial confinement, and growth-rate measurement, so the reversal cannot be attributed to a single environmental feature. Nevertheless, context dependence is consistent with a specialist state favored by the RQ selection regime. Second, the detected mutations in the endpoint clones did not overlap: RQ70 carries *polA* R821C, a 13,484-bp deletion spanning 16 flagellar and chemotaxis genes, and an insertion disrupting FHr *traA*, whereas MG70 carries *rpoA* N294H (**Figure 6B**). The phenotypic contribution of the *traA*-disrupting FHr insertion remains unresolved. Variations on the flagellar and chemotactic deletion have been described in the ALEdb, while *polA* R821C and *rpoA* N294H have not.^53^

RQ70’s broader phenotype reinforces its interpretation as a context-dependent specialist. Across the first phase of longitudinal RQ measurements, maximum growth rate and lag time were moderately positively associated (*r* = 0.41; **Figure 4D**), indicating that longer lag accompanied the evolutionary trajectory rather than appearing only at its endpoint. RQ70’s lag time was 3.03 h, compared with 1.42 h for MG70 and 1.12 h for the ancestor. It also showed less robust outgrowth at intermediate-to-high sub-MIC concentrations of gentamicin, kanamycin, and chloramphenicol, but tolerated the composite plasmid–kanamycin challenge better than either comparator (**Supplementary Figures 8 and 14–17**). These shifts are consistent with prior work showing that microbial fitness is multidimensional and that faster growth can trade off against adaptation time, yield, or stress performance.^38–40,54,55^ These apparent deficiencies provide a plausible explanation for why a phenotype resembling RQ70 has not commonly emerged from conventional adaptive laboratory evolution.

Together, these results support a general principle: an apparent physiological plateau can be shaped by the structure of selection under which it is measured. By engineering the mapping from phenotype to fitness, amplified selection may expose both physiological capacities and evolutionary paths that composite selection does not readily access. Existing synthetic selections have enriched high metabolite producers, isolated hypersecretory *E. coli* phenotypes, and stabilized costly biosynthetic activity during long-term cultivation.^43,56,57^ RQ shows that the same design principle can be used not only to optimize an engineered output but also to interrogate an apparent physiological ceiling.

## Methods

### Strains and growth conditions

All experiments were performed using either *Escherichia coli* Top10F’ or *Escherichia coli* K-12 MG1655 carrying the FHr plasmid (MG1655+FHr). Cultures were grown at 37°C with shaking at 225 rpm unless otherwise specified. Liquid cultures were grown in Apex LB Broth (Miller) Mix (Cat No. 11-120, Lot No. R7A95240). Solid media consisted of Apex LB Agar (Miller) Mix (Cat No. 11-122, Lot No. S912R0E). Antibiotics were used at the following concentrations unless otherwise noted: chloramphenicol (Cm), 25 µg/mL; carbenicillin (Carb), 100 µg/mL. Anhydrotetracycline (aTc) was used at concentrations ranging from 0.01 to 100 ng/mL to induce circuit activity, as specified for individual experiments.

### Construction of the Red Queen (RQ) gene circuit

The Red Queen (RQ) circuit consists of two plasmids: pCas9 and pTarget. pCas9 carries the gene for a catalytically active Cas9 endonuclease fused to an ssrA degradation tag and expressed from a tetracycline-inducible promoter (pTet). pCas9 contains a low-copy-number SC101 replication origin and confers chloramphenicol resistance.

pTarget carries a guide RNA (gRNA) driven by pTet that targets a non-coding region on the same plasmid. pTarget also encodes a constitutively expressed β-lactamase (AmpR), which is essential for survival in the presence of carbenicillin, and uses a high-copy-number pUC origin of replication.

Upon aTc induction, Cas9–gRNA complexes cleave pTarget, reducing AmpR copy number. Because Cas9 accumulation depends on dilution through cell division, the circuit imposes growth-rate–dependent survival: slow-growing cells accumulate Cas9 and lose pTarget, while fast-growing cells dilute Cas9 sufficiently to maintain AmpR expression and survive antibiotic selection. Control constructs included pTarget plasmids encoding a mismatched, non-targeting gRNA.

### Assessment of growth-rate-dependent survival on solid and liquid media

RQ-carrying strains were streaked onto LB agar plates containing Cm and Carb, with or without 100 ng/mL aTc. Plates were incubated for 24 h at 37 °C. Individual colonies were then inoculated into LB liquid media containing either Cm alone or Cm + Carb and grown for 16 h. Optical density at 600 nm (OD600) was measured to assess survival and outgrowth. Experiments were performed with targeting and non-targeting gRNAs and with aTc present or absent.

### Dilution-based growth escape assays

To test whether fast-growing cells could escape RQ-mediated suppression, dilution-based growth experiments were performed. Cultures were grown in LB containing Cm and Carb, with or without aTc induction. In the exponential-growth condition, cultures were diluted 1,000-fold into fresh media every 12 hours to maintain high effective growth rates. In the slow-growth control condition, cultures were grown for 24 hours without dilution before being diluted 1,000-fold. OD600 measurements were taken every 12 hours. Six biological replicates were used for each condition.

### The selection pipeline to evolve Top10F’

RQ cells (Top10F’) carrying either a targeting (sp) or non-targeting (NT) gRNA, and naïve Top10F’ cells, were streaked onto LB agar plates containing 25 µg/mL Cm + 100 µg/mL Carb (for RQ lineages) or no antibiotics (for naïve lineages). After overnight incubation (16 h, 37 °C), four colonies of sp RQ cells, two colonies of NT RQ cells, and three colonies of naïve cells were selected as biological replicates and inoculated into 1 mL of the appropriate medium in a 2 mL deep-well plate for overnight growth. Overnight cultures were then diluted 10,000-fold into four media conditions: LB + 25 µg/mL Cm + 100 µg/mL Carb supplemented with 0, 0.1, 1, or 10 ng/mL aTc (RQ lineages), or antibiotic-free LB (naïve lineages). Cultures were passaged daily by 10,000-fold dilution for 43 days.

On Day 43, the two fastest growing sp RQ lineages (both evolved in 0.1 ng/mL aTc), one NT RQ lineage (from 0.1 ng/mL aTc), and all three naïve lineages were selected for continued evolution. The selected RQ lineages were passaged under more stringent selection — 100,000-fold daily dilution into LB + Cm + Carb + 1 ng/mL aTc — for an additional 57 days, while the naïve lineages continued in antibiotic-free LB at the same 100,000-fold dilution. Growth rates at Days 1, 43, and 100 were measured in a plate reader by diluting overnight cultures 1,000-fold into 200 µL of LB in a 96-well plate, sealing with 50 µL mineral oil to prevent evaporation, and recording OD600 every 10 min for at least 12 h with orbital shaking.

### Short-term aTc titration

MG1655+FHr cells carrying the RQ circuit were streaked onto LB agar containing 25 µg/mL Cm and 100 µg/mL Carb and incubated for 16 h at 37 °C. Four colonies were picked as biological replicates and grown overnight in 1 mL LB + Cm + Carb in 2 mL deep-well plates. Overnight cultures were passaged daily into LB + Cm + Carb supplemented with aTc at 0, 0.1, 0.2, 0.3, 0.6, 1, 2, 3, 6, and 10 ng/mL, at daily dilution factors of either 1:10,000 or 1:100,000, for two consecutive days. On Day 3, cultures were diluted 1,000-fold into 200 µL LB in a 96-well plate, sealed with SealPlate film, and OD600 was recorded every 10 min for at least 12 h at 37 °C with orbital shaking in a Tecan Infinite 200 PRO. Maximum specific growth rate was determined by fitting Equation S9 to each trace. Three technical replicates were measured per biological replicate.

### The selection pipeline to evolve MG1655+FHr

The long-term evolution experiment was conducted over 70 days total, according to the schedule described in **Supplementary File 1**. RQ populations evolved in LB containing Cm and Carb under gradually increasing aTc concentrations. Control populations evolved in LB without antibiotics or induction.

Lineages were propagated in 96-well deep well plates with 1 mL of the appropriate media in each well. Each day, cells from overnight were passaged into an “activation” plate by diluting 100-fold (before Day 29) or 1,000-fold (after Day 29) into LB + Cm + Carb (for RQ lineages) or LB (for negative controls). The activation plates were incubated for 2 hours. After incubation, the lineages were diluted 100-fold into the “treatment” plate with LB + Cm + Carb + aTc (for RQ lineages) or LB (for negative controls). Approximately every other day, the lineages were also diluted 100-fold from the activation plate into 96-well plates, sealed with SealPlate film, and OD600 was recorded every 10 min for at least 12 h at 37 °C with orbital shaking in a Tecan Infinite 200 PRO.

A lineage selection strategy was employed in which the fastest-growing lineages were propagated, while slower-growing lineages were discarded. This selection occurred approximately every 14 days. Growth rates were periodically measured using plate reader growth curves by diluting overnight cultures 1,000-fold into 200 µL of LB in a 96-well plate, sealing with Excel Scientific SealPlate film, and recording OD600 every 10 min for at least 12 h with orbital shaking. Growth rate was quantified as the maximum specific growth rate of a logistic-with-lag fit to each trace. Overly strong induction (>10 ng/mL aTc) was avoided due to circuit instability and growth inhibition.

### Circuit curing and shake-flask growth measurements

To assess whether growth improvements persisted after removal of the RQ circuit, evolved strains were cured of plasmids by serial passaging without antibiotic selection. Loss of plasmids was confirmed by antibiotic sensitivity.

RQ-circuit-cured RQ70 colonies, MG70 colonies, and ancestral MG1655+FHr colonies were picked from streak plates into overnight cultures. Following million-fold dilution the following morning, they were grown in 20 mL LB cultures in baffled shake flasks at 37 °C. Growth of RQ70, MG70, and ancestral MG1655+FHr shake-flask cultures was quantified by viable colony counts. Cultures were sampled hourly from t = 0–10 h, and each timepoint was plated in replicate; replicate counts within a biological replicate were averaged to give a single CFU/mL trajectory per flask (RQ70, n = 12; MG70, n = 3; MG1655+FHr, n = 3 biological replicates).

Instantaneous growth rates were estimated by a sliding-window log-linear regression. For every candidate start time t₀, counts at t₀ through t₀ + 3 h (four consecutive hourly samples) were log-transformed and ln(CFU/mL) was regressed against time by ordinary least squares; the slope of this fit was taken as the growth rate μ (h⁻¹) for that window. Windows containing fewer than three sampled timepoints or any non-positive count were discarded. This yielded a rolling 3-h growth-rate series per biological replicate.

The maximum growth rate *μ_max_* for each biological replicate was defined as the largest slope across all valid windows, and the corresponding doubling time was calculated as *ln*2/*μ_max_*. Conditions were compared by one-way ANOVA with Tukey’s HSD post-hoc test; Levene’s test and a Kruskal–Wallis test were run alongside to confirm that the conclusion was not contingent on the equal-variance or normality assumptions.

### Turbidostat continuous culture

Continuous culture was performed in a custom microplate-based turbidostat operating at 37 °C. Overnight cultures of RQ70, MG70, and ancestral MG1655+FHr were diluted 1,000-fold into 1 mL of LB per well and grown to the target optical density, after which the instrument maintained density by automated dilution with fresh LB. No antibiotics or inducers were present in any turbidostat culture. Four OD setpoints were tested (0.6, 0.7, 0.8, and 0.9), with eight biological replicate wells per strain per setpoint. Optical density was recorded every 20 minutes and dilution events were logged throughout.

Instantaneous specific growth rate was calculated from the dilution record as the rate of medium addition required to hold OD constant. Steady-state growth rate was taken as the mean of the instantaneous rate over the window t = 5–10 h, after visual confirmation that the time series had plateaued (**Supplementary Figure 9A**).

### Mother machine measurements

Mother machine devices were fabricated as previously described.^58^ Overnight cultures of RQ70, MG70, and ancestor strains were diluted 1:100 and grown for 3 h at 37 °C before being loaded in separate channels of a mother machine. The device was next spun at 4696 x g for 5 min before LB + 0.2 mg/mL Pluronic F-127––to avoid cell adhesion to the device––was flowed through the channels. The device was then mounted on a Nikon Eclipse Ti2 inverted microscope stage held at 37 °C, and cells were allowed to recover for 3 h. Next, single cells were imaged in phase contrast every 5 min for 16 h, using a 50 ms exposure time. DeLTA software was used on-the-fly to segment cells and extract cell features (i.e., cell area, perimeter, width, and length) from phase contrast images.^59,60^ The extracted cell area was used to calculate growth rate (in ℎ^−1^) following the below equation:

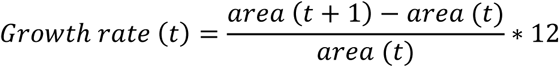

At any time point, data from cells with a growth rate above 3.6 ℎ^−1^ or below -2.4 ℎ^−1^ as well as cells with areas too small to represent a true cell were filtered out as they represent either cell division events or non-physiological data points generated by segmentation errors.

### Agar pad microscopy

Cells were inoculated from a single colony into 3 mL of LB and incubated at 37 C, 225 rpm overnight. The overnight culture was then diluted 1:100 into 3 mL of LB. When cells reached OD600 ∼0.4, cells were diluted 1,000-fold and 2.25 μL of this dilution was spotted onto an agarose pad. After drying for 5 min, the agarose pad was inverted into a coverslip dish, sealed with parafilm, and then secured into a Tokai Hit stage-top incubator on the microscope stage (Keyence BZ-X810). The incubator was set to 37 °C. Using an automated stage, Z-stacks of phase contrast images were taken every 3 min in multiple locations with a 40x phase contrast objective lens.

Images were cropped to remove debris from the image, and in-focus images were then selected using a custom routine. This involved calculating the Tenengrad metric around a manually selected region of interest for the first 60 frames, followed by calculating the metric for the entire image in the remaining frames.

Cells were segmented using CellProfiler 4.2.6 and Omnipose.^61,62^ Cell tracking was performed using the overlap method in CellProfiler. Neighbors for each cell were identified by expanding each segmented cell by 3 pixels (0.57 μm). Lineage tracking correction was then performed using TrackRefiner4, with default settings.^63^ Division threshold length was defined as the final length of a cell the frame prior to its division. Cell radius was defined as half the minor-axis of the ellipse fitted to a segmented cell. Single-cell growth rates were computed over the cell cycle through ordinary least-squares regression of the log of the cell length versus time.

Multiple filtering steps were performed prior to analysis. *E. coli* MG1655 has a radius of around 0.5 μm, so objects with a measured radius greater or equal to 1 μm (approximately 2x the known radius of *E. coli* MG16555) were filtered out, since an *E. coli* cell with a larger radius would be caused by a merging error in the cell segmentation algorithm. Cells with a growth rate less than or equal to 0 μm/hour were also filtered out, as these cells would be considered non-growing or dying. Cells whose fitted growth rate had a correlation coefficient (R^2^) less than 0.95, or less than four data points were also filtered out to reduce chances of measuring an artificially enhanced growth rate due to cell tracking errors.

### Growth in M9 minimal medium

M9 minimal medium was prepared as 1× M9 salts, 2 mM MgSO4, 0.1 mM CaCl2, and 1% (w/v) thiamine with 0.4% (w/v) glucose as the sole carbon source. Overnight cultures of RQ70, MG70, and ancestral MG grown in M9 were diluted 40-fold into 200 µL M9 + 0.4% glucose in a 96-well plate, with 12 replicate wells per strain. Plates were sealed with SealPlate film and incubated at 37°C with orbital shaking.

### Plasmid carriage under kanamycin selection

To assess growth during plasmid carriage under kanamycin selection, RQ70, MG70, and ancestral MG1655+FHr were transformed by heat shock transformation with either a medium-copy p15A-origin plasmid or the high-copy pADEPT plasmid, both conferring kanamycin resistance. Transformants were selected on LB agar containing 50 µg/mL kanamycin.

Growth was measured in M9 + 0.4% glucose as described above, with six replicate wells per strain–plasmid combination supplemented with 50 µg/mL kanamycin, alongside twelve plasmid-free wells of the same strain in M9 without kanamycin.

### Antibiotic responses

Growth in response to gentamicin, kanamycin, and chloramphenicol was assessed by broth microdilution in 96-well plates. Each plate contained a two-fold dilution series from 64 to 0 µg/mL across columns 2–11, with column 12 containing LB alone as a blank. Rows B–E were inoculated with the strain of interest (n = 4 wells) and rows F–G with the ancestral MG1655+FHr control (n = 2 wells per plate, pooled across plates to n = 4).

Overnight cultures were diluted 1,000-fold into 200 µL LB per well containing the appropriate drug concentration. Plates were sealed with 50 µL of mineral oil and growth curves were measured as described previously. Apparent MIC within the assay window was defined as the lowest concentration at which no growth was detected, where growth was defined as blank-subtracted OD600 exceeding 0.05.

### Whole-genome sequencing and mutation analysis

Overnight cultures of the RQ70 clone, the MG70 evolved control, seven single clones isolated from the RQ70 population, a pooled population (“Meta”) sample, and the ancestral MG1655+FHr strain were preserved in Zymo 1× DNA/RNA Shield and submitted to Plasmidsaurus for genomic DNA extraction. Library preparation and sequencing were performed by Plasmidsaurus. Amplification-free long-read libraries were constructed using Oxford Nanopore v14 chemistry with sequence-independent fragmentation of input genomic DNA by tagmentation, and sequenced using a primer-free protocol on R10.4.1 flow cells. Reads were basecalled with Dorado v4.3 in super-accurate mode with default Q10 quality filtering, then filtered with Filtlong v0.2.1 to remove reads <300 bp and retain the best 95% by quality. Genome size was estimated with Lrge v0.2.1, and multiple subsampled read sets were assembled with Autocycler using Flye v2.9.6, hifiasm, and Plassembler v1.8.0; low-depth and small contigs were removed and the resulting assemblies compressed, clustered, trimmed, and resolved by Autocycler to a single consensus assembly, which was rotated to an optimal start position with dnaapler. Assemblies were annotated with Bakta v1.11, assessed for completeness and contamination with CheckM v1.2.2, and replicon identity confirmed with Sourmash v4.9.4 against GTDB rs226, RefSeq plasmid, and phage databases. Assembly consensus accuracy for this service is typically Q50– Q60.

Mutations were identified by whole-genome alignment of de novo assemblies. Each evolved assembly was aligned against the ancestral Day 0 MG1655+FHr assembly using minimap2 with the asm5 preset for assembly-to-assembly comparison at low divergence. Substitutions, insertions, and deletions were extracted by parsing the cs difference string of each alignment. Two comparisons were performed against the same ancestral reference: (i) the evolved MG70 control and the plasmid-cured RQ70 clone; and (ii) seven single clones isolated from the RQ70 population together with a pooled population (“Meta”) sample.

#### Modeling RQ circuit dynamics

The RQ model consists of three ordinary differential equations (ODEs), each of which describes the dynamics of cell population (*N*), average per-cell pTarget copy number (*P*) and intracellular endonuclease Cas9 concentration (*E*).

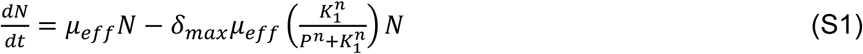

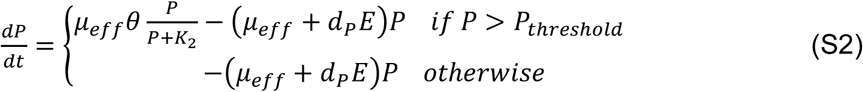

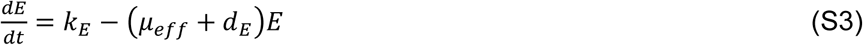

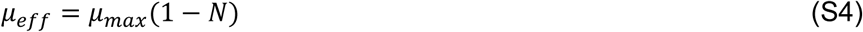

The model has ten parameters: the maximum cell growth rate (*μ_max_*), the maximum RQ circuit-mediated cell death rate (*δ_max_*), the pTarget copy number-mediated cell rescue constant (*K*_1_), the Hill coefficient for the cell death rate (*n*), the pTarget replication rate (*θ*), the pTarget replication Michaelis constant (*K*_2_), the Cas9-mediated plasmid digestion rate (*d_P_*), the threshold of pTarget copy number for replication (*P_t_*_ℎ*res*ℎ*old*_), the maximum Cas9 expression rate (*k_E_*), and the Cas9 degradation rate (*d_E_*). The model also factors in the growth rate-dependent dilution of the intracellular pTarget and endonuclease concentrations (**Equations S2 and S3**).

To model population-level adaptation under selection with the RQ circuit, we added to the model as follows:

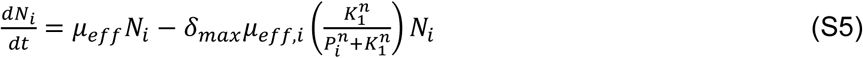

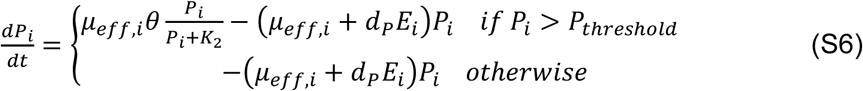

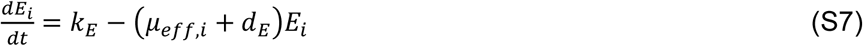

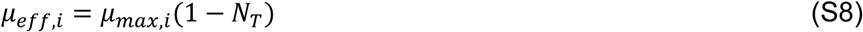

The subscript *i* indexes the member of the population and the total population size *N_T_* = ∑ *N_i_*. See **Supplementary Table 1** and **Supplementary Table 2** for parameter values used in simulations.

Population-adaptation simulations were implemented in Python using scipy.integrate.solve_ivp (RK45). Each replicate initialized 5,000 lineages with maximum growth rates drawn independently from a normal distribution with mean *μ*₀ = 1.0 and s.d. *σ*₀ = 1 × 10⁻⁴, an initial coefficient of variation of 0.01%. Lineages were propagated through 25 sequential passages of 24 time units. Within each passage all lineages competed for a shared normalized carrying capacity (*ΣN* = 1) and were seeded at *N* = 1/*V* with *V* = 1 × 10⁹, so *N* = 1/*V* corresponds to a single cell at capacity. Adaptive evolution integrated logistic competition in N alone; amplified selection additionally integrated the plasmid (P) and Cas9 (E) states of the RQ circuit at static induction *k_E_* = 1.0 with *δ_max_* = 1, *K*₁ = 0.7, *n* = 3, *θ* = 1, *K*₂ = 0.1, *d_P_* = 1, *P_t_*_ℎ*res*ℎ*old*_ = 0.1, and *d_E_* = 0.05.

At the end of each passage, lineages below 1/V were discarded and each surviving lineage i contributed descendants drawn independently from a Poisson distribution with mean 5,000 × *N*ᵢ/*ΣN*. Each descendant’s maximum growth rate was drawn from a normal distribution centered on its parent’s value with s.d. equal to 1% of that value. A replicate was recorded as extinct when total end-of-passage abundance fell below 1 × 10⁻⁴ of carrying capacity or when fewer than two lineages remained; thereafter mean growth rate was held at its last value and survival fraction set to zero.

### Logistic model of cell growth with lag

To determine maximum growth rates and lag times from growth curves, the following piecewise equation was fit to the growth curve traces:

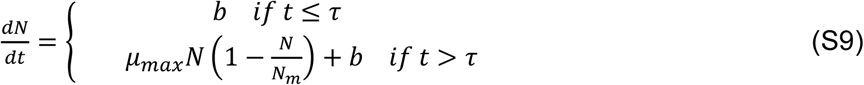

where *b* is the baseline drift of non-exponential growth, *τ* is the lag time, and *N_m_* is the carrying capacity.

## Acknowledgments

The authors would like to thank Amy Schmid, Michael Lynch, and Ophelia Venturelli for insightful comments and suggestions. This work was partially supported by the National Science Foundation (G.S.H.: DGE 2139754) and the National Institutes of Health (G.S.H: 1T32GM144291; L.Y.: R01GM098642 and R01EB031869).

## Author contributions

G.S.H., H.-I.S., and L.Y. devised the project. G.S.H., H.-I.S., R.M., and L.Y. formulated theoretical framework. G.S.H. and H.-I.S. performed circuit characterization experiments. G.S.H., H.-I.S., Z.Z., K.L., X.C., A.Y., J.M.Q., C.V., Q.M., H.M., I.S., and A.R.S. contributed to evolution and characterization of *E. coli* strains. E.J.C., M.J.D., and L.Y. supervised the work and acquired funding. G.S.H., H.-I.S., and L.Y. wrote the manuscript with input from all authors.

## Data availability

Data is available in the main text and the supporting information, on request from the corresponding author, and available directly with code at https://github.com/ghamrick34/RQ.

## Supplementary information

**Supplementary Figure 1.**
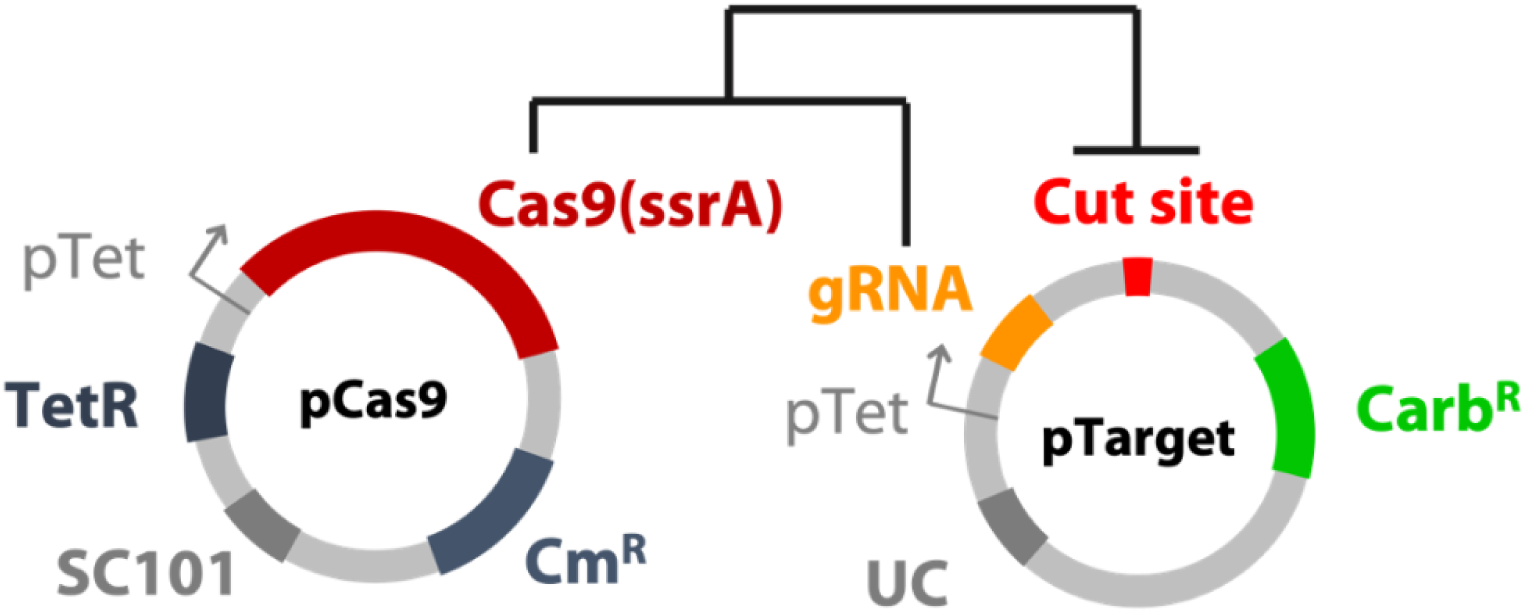
Red Queen circuit schematic. The circuit consists of two plasmids, pCas9 and pTarget. pCas9 encodes a Cas9 protein tagged with a protease degron sequence under the tetracycline promoter (pTet) and features a low copy number replication origin to reduce Cas9-induced cytotoxicity. pTarget expresses a guide RNA (gRNA) controlled by pTet. Upon induction with anhydrotetracycline (aTc), the Cas9-gRNA complex targets and cleaves a non-coding region on pTarget. pTarget also carries the carbenicillin resistance (Carb^R^) gene essential for the cell survival in the presence of the antibiotic. pCas9 confers resistance to chloramphenicol (Cm^R^).

**Supplementary Figure 2.**
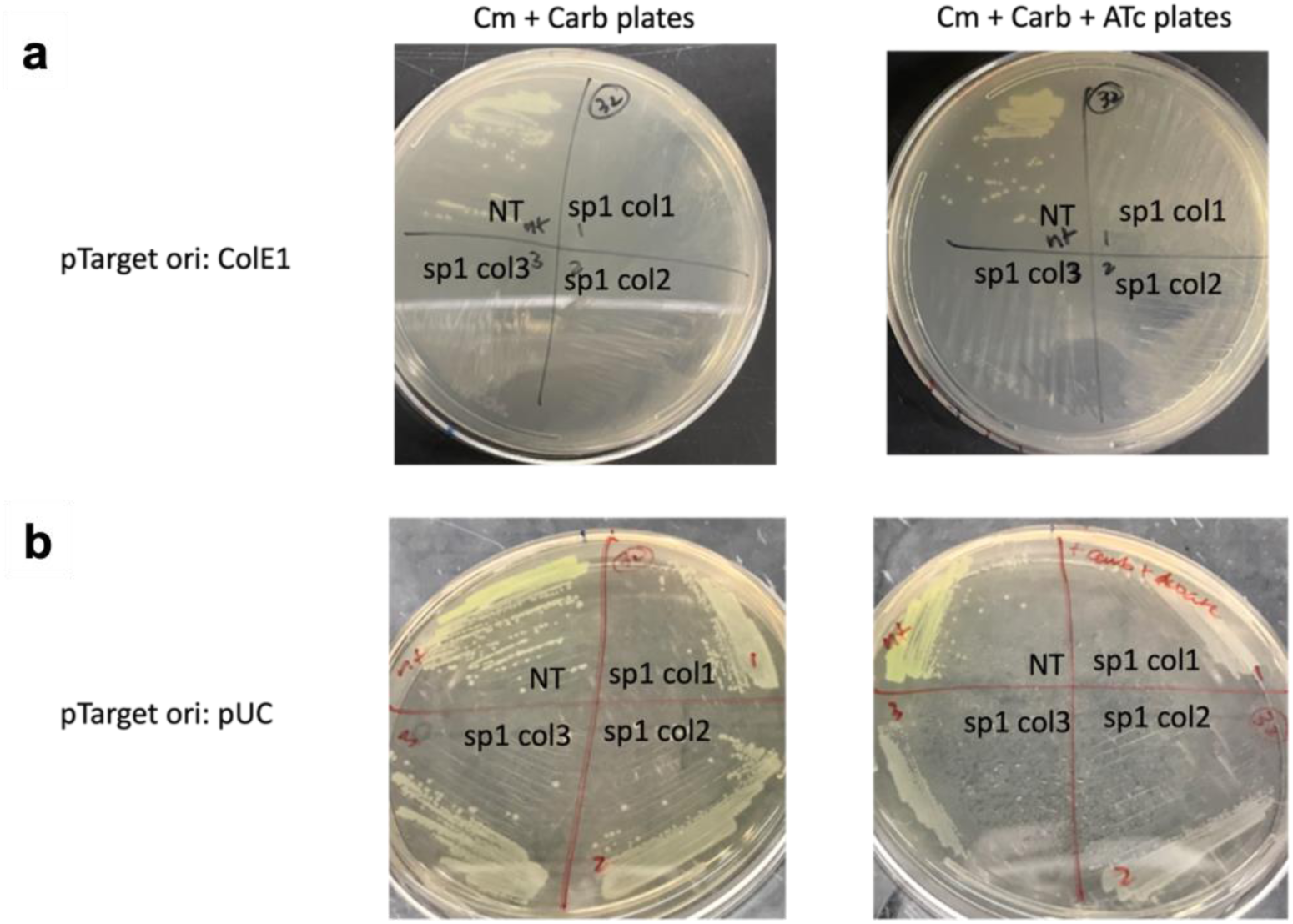
LB agar plates streaked with RQ cells: plasmid backbone optimization. **(a)** An earlier iteration of the RQ circuit using a medium-copy ColE1 replication origin for pTarget did not grow on LB agar plates containing 25 µg/mL Cm and 100 µg/mL Carb (no aTc), indicating that basal Cas9 cytotoxicity was sufficient to prevent cell growth even with the circuit OFF. Three colonies of targeting (sp) RQ cells were streaked per plate but none grew. Non-targeting (NT) RQ cells grew in both the presence and absence of aTc, confirming that the failure of sp cells was Cas9-dependent. **(b)** The SC101/pUC configuration used throughout this study grew robustly on LB agar plates supplemented with 25 µg/mL Cm, 100 µg/mL Carb, and either 0 or 100 ng/mL aTc. Plates shown are from t = 24 h of the experiment presented in Figure 2. RQ cells carrying the targeting spacer had slightly smaller colonies on aTc-containing plates, consistent with growth-rate–dependent suppression.

**Supplementary Figure 3.**
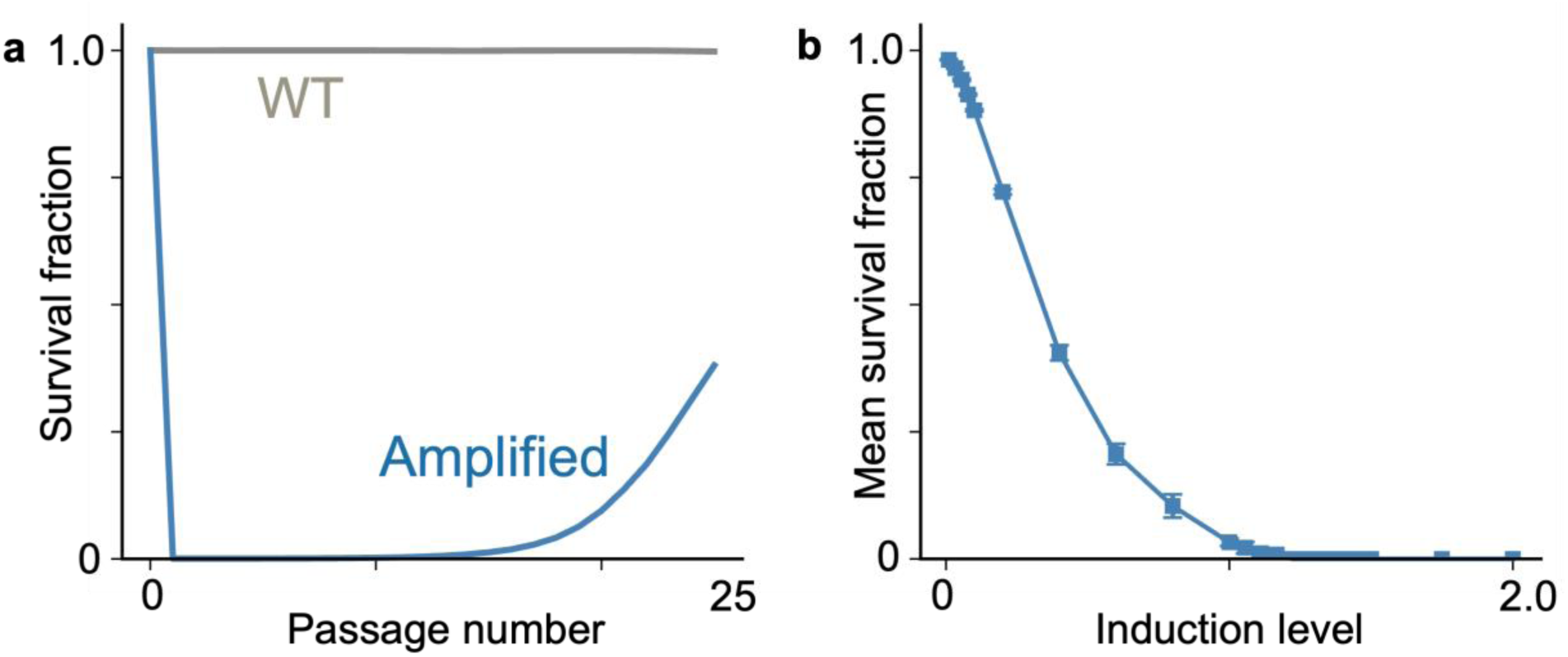
Survival dynamics under amplified selection. **(a)** Per-passage survival fraction for WT (gray) and amplified selection at *k_E_* = 1.0 (blue). The dashed red line marks the extinction threshold (0.01%). **(b)** Mean survival fraction per passage across the *k_E_* sweep. Error bars indicate s.d. across replicates.

**Supplementary Figure 4.**
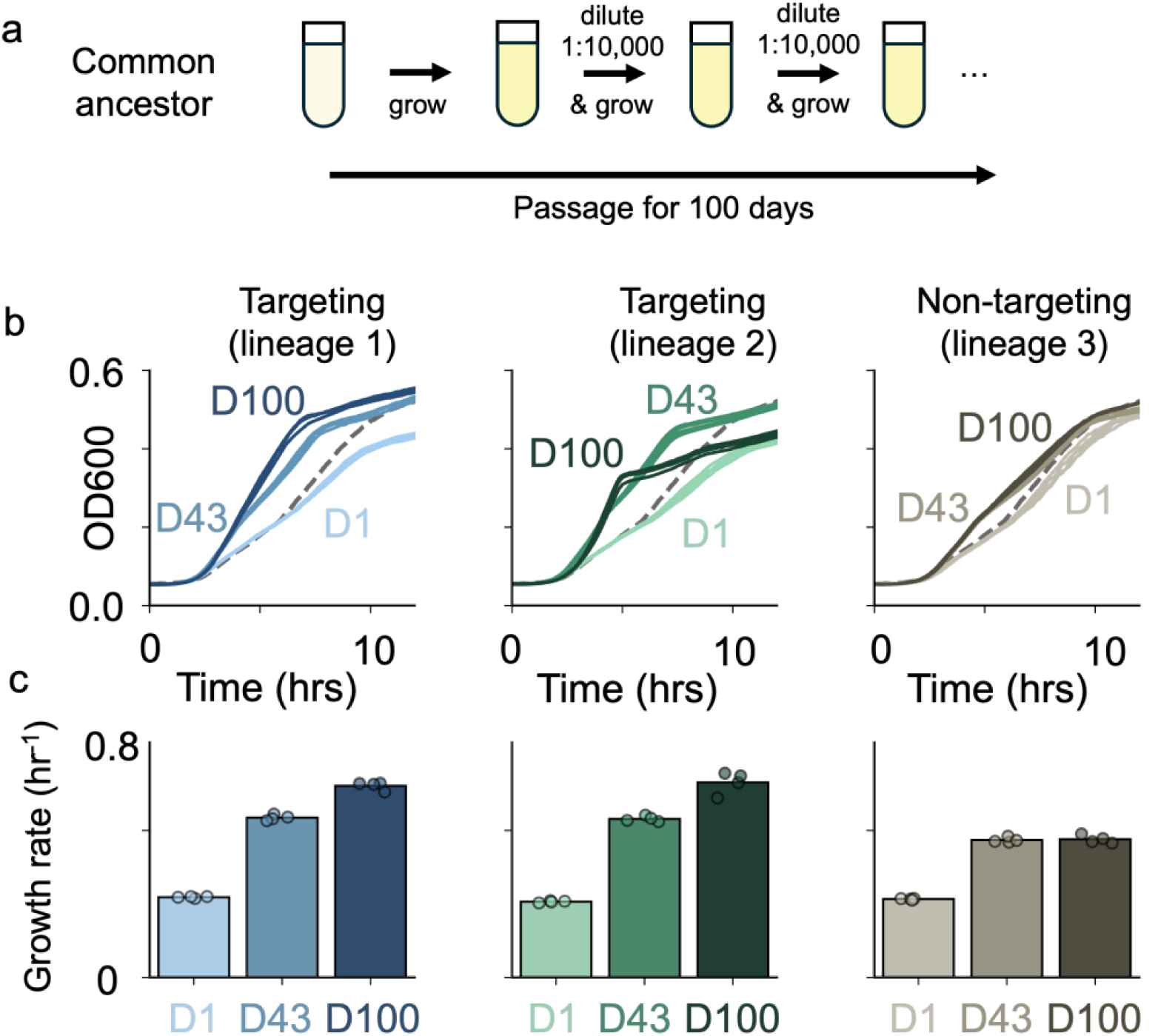
Amplified selection accelerates *E. coli* Top10F’ growth rate over 100 days of serial passaging. **(a)** Experimental protocol. Four lineages of RQ cells carrying a targeting gRNA (sp), two lineages carrying a non-targeting gRNA (NT), and three naïve *E. coli* Top10F’ lineages were passaged daily for 100 days. For the first 43 days, cultures were diluted 10,000-fold into LB + Cm + Carb supplemented with 0, 0.1, 1, or 10 ng/mL aTc. On Day 43, the two fastest-growing targeting lineages and one non-targeting lineage (all evolved in 0.1 ng/mL aTc) were selected for continued evolution under 100,000-fold daily dilution with 1 ng/mL aTc for the remaining 57 days. **(b)** Growth curves of representative ancestor (D1) and evolved (D43, D100) RQ cells for targeting lineage 1 (blue), targeting lineage 3 (green), and non-targeting lineage 2 (gray). Darker shading corresponds to more evolved strains. Dashed gray line shows the mean Day 1 naïve ancestor growth curve. Each curve represents four technical replicates. **(c)** Maximum growth rates of the three RQ lineages from (B) across Days 1, 43, and 100. By Day 100, targeting lineages achieved 1.95- and 1.86-fold improvements in growth rate, while the non-targeting lineage achieved a 1.16-fold improvement. Individual data points represent four technical replicates.

**Supplementary Figure 5.**
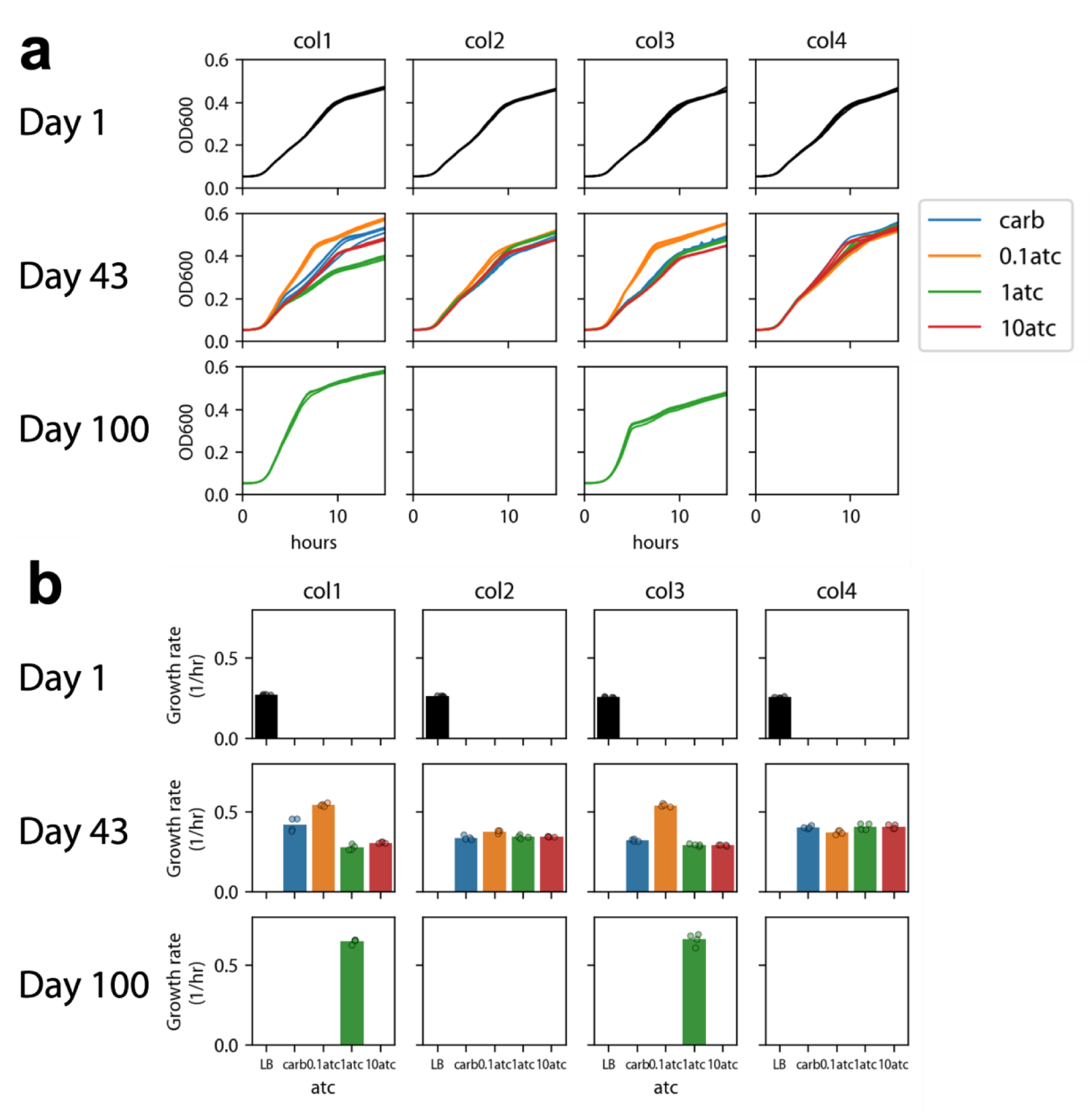
Growth curves of the 100-day LTAEE RQ cells carrying a targeting spacer (Top10F’) Growth curves **(a)** and maximum growth rates **(b)** of all four biological replicates (col1–col4) of RQ cells carrying the targeting (sp) gRNA from Days 1, 43, and 100 of the Top10F’ LTAEE. On Day 1, each colony was inoculated into four media types (LB + 25 µg/mL Cm + 100 µg/mL Carb supplemented with 0, 0.1, 1, or 10 ng/mL aTc). After 43 days of passaging at 10,000-fold daily dilution, col1 and col3 evolved in 0.1 ng/mL aTc showed the greatest growth-rate improvement and were selected for an additional 57 days of evolution under 100,000-fold daily dilution with 1 ng/mL aTc. These strains improved further by Day 100. Each growth curve represents four technical replicates; bars in **(b)** show mean growth rates across replicates with individual data points shown as circles.

**Supplementary Figure 6.**
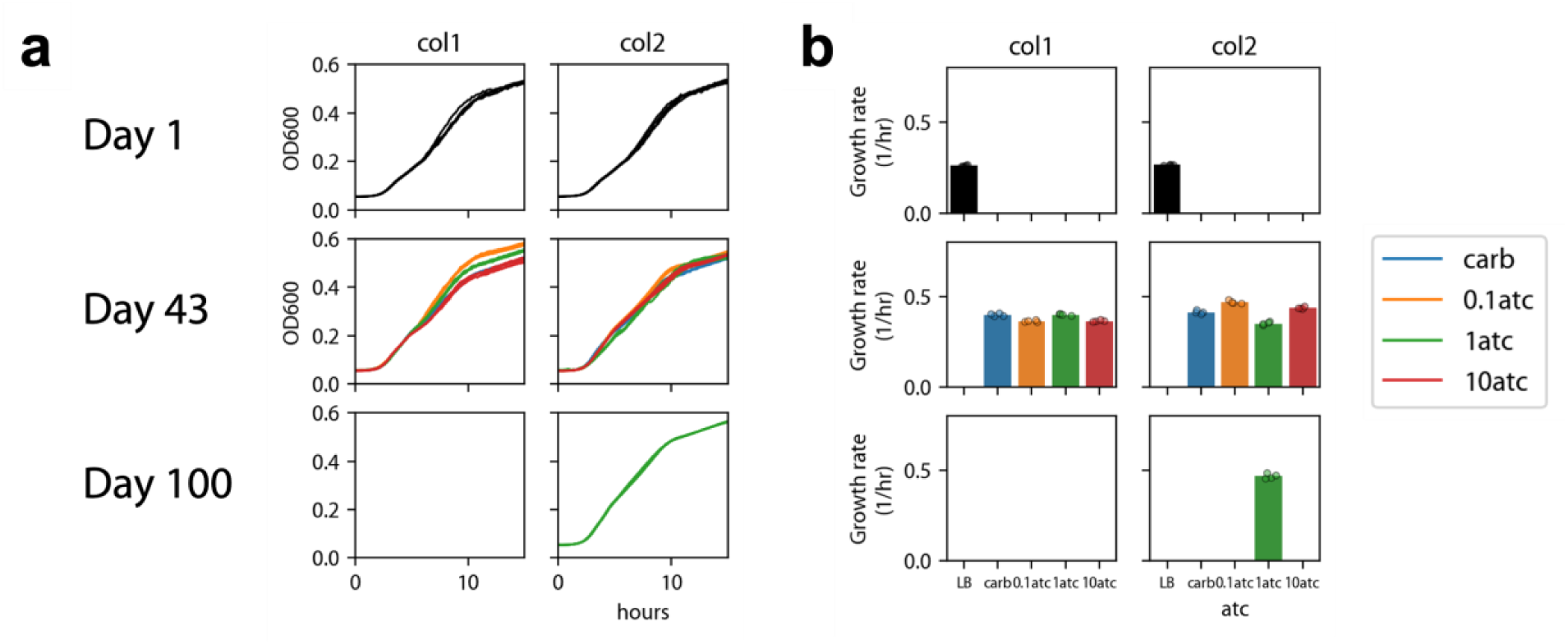
Growth curves of the 100-day LTAEE RQ cells carrying a non-targeting spacer (Top10F’) Growth curves **(a)** and maximum growth rates **(b)** of both biological replicates (col1, col2) of RQ cells carrying the non-targeting (NT) gRNA from Days 1, 43, and 100 of the Top10F’ LTAEE. On Day 1, each colony was inoculated into four media types (LB + 25 µg/mL Cm + 100 µg/mL Carb supplemented with 0, 0.1, 1, or 10 ng/mL aTc). On Day 43, col2 (evolved in 0.1 ng/mL aTc) was selected for an additional 57 days of evolution under 100,000-fold daily dilution with 1 ng/mL aTc, producing a 1.16-fold growth-rate increase by Day 100. Each growth curve represents four technical replicates; bars in **(b)** show mean growth rates across replicates with individual data points shown as circles.

**Supplementary Figure 7.**
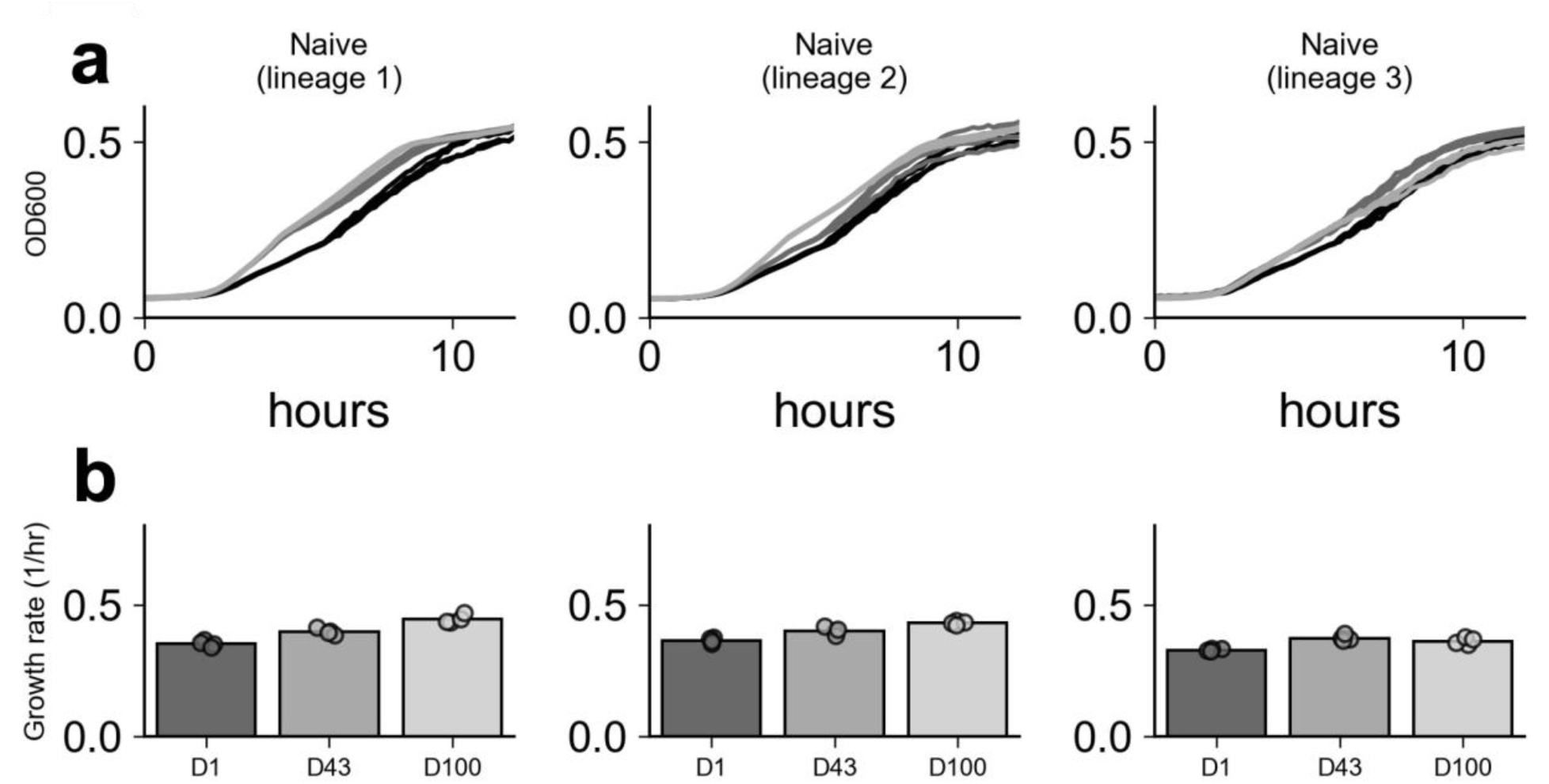
Growth curves of the 100-day LTAEE naïve cells (Top10F’) (A) Raw growth curves of the ancestor (D1) and adapted (D43, D100) Top10F’ cells from three lineages. (B) Maximum growth rates of the three lineages at Days 1, 43, and 100. Naïve lineages, passaged in antibiotic-free LB, accelerated 1.10- to 1.27-fold over the 100-day experiment. Bars show mean growth rates across four technical replicates with individual data points shown as circles.

**Supplementary Figure 8.**
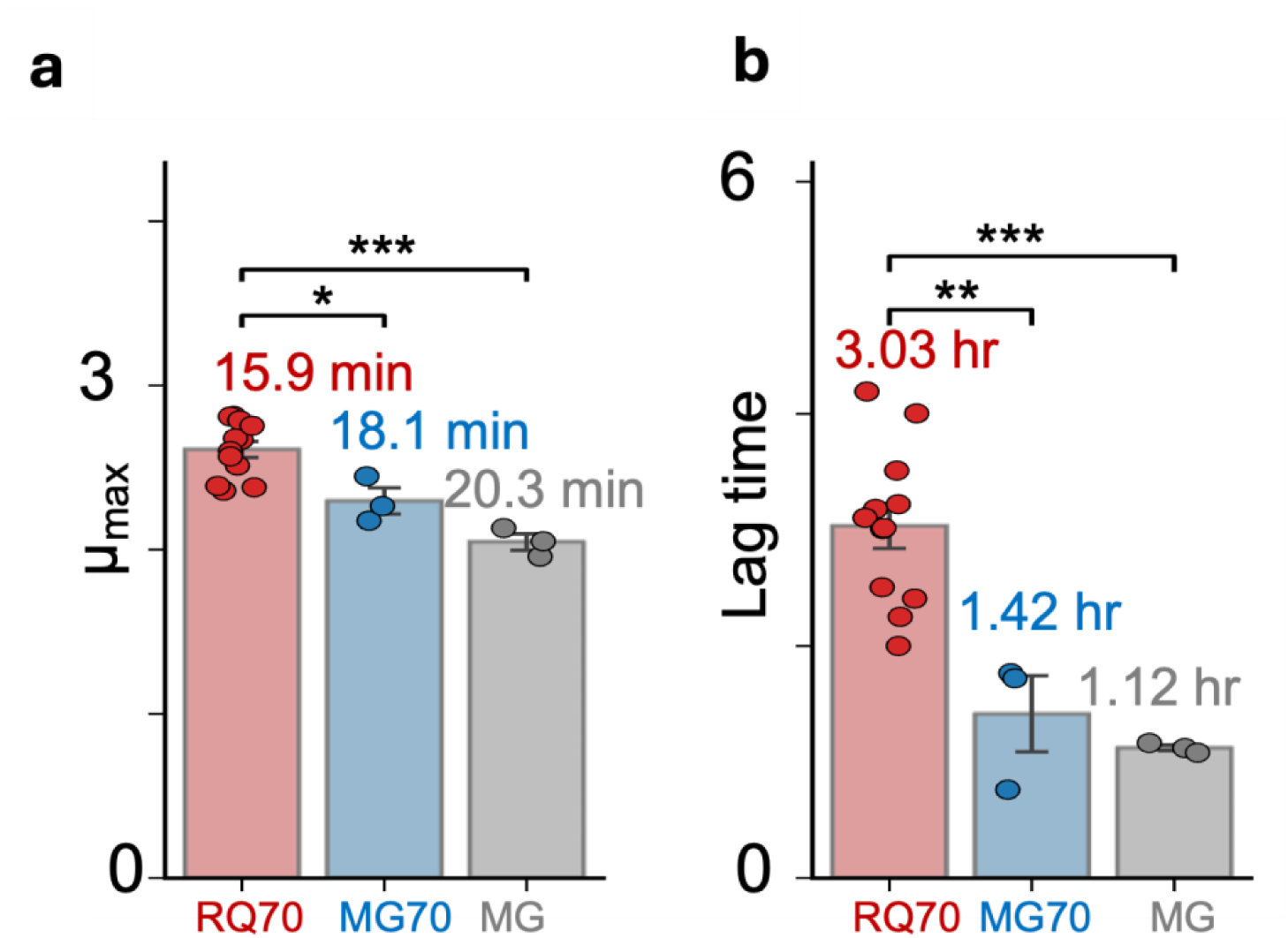
RQ70 achieves the highest maximum growth rate but the longest lag time in shake-flask culture. Maximum specific growth rate (A) and lag time (B) for RQ-circuit-cured RQ70 (red), the Day 70 evolved control MG70 (blue), and the ancestral MG1655+FHr strain (gray), measured by CFU counting in 20 mL baffled shake-flask cultures in LB at 37 °C without antibiotics or inducers. Growth rates were calculated using a 3-hour rolling window; doubling times (shown above each bar) were computed as 60·ln(2)/µ.

**Supplementary Figure 9.**
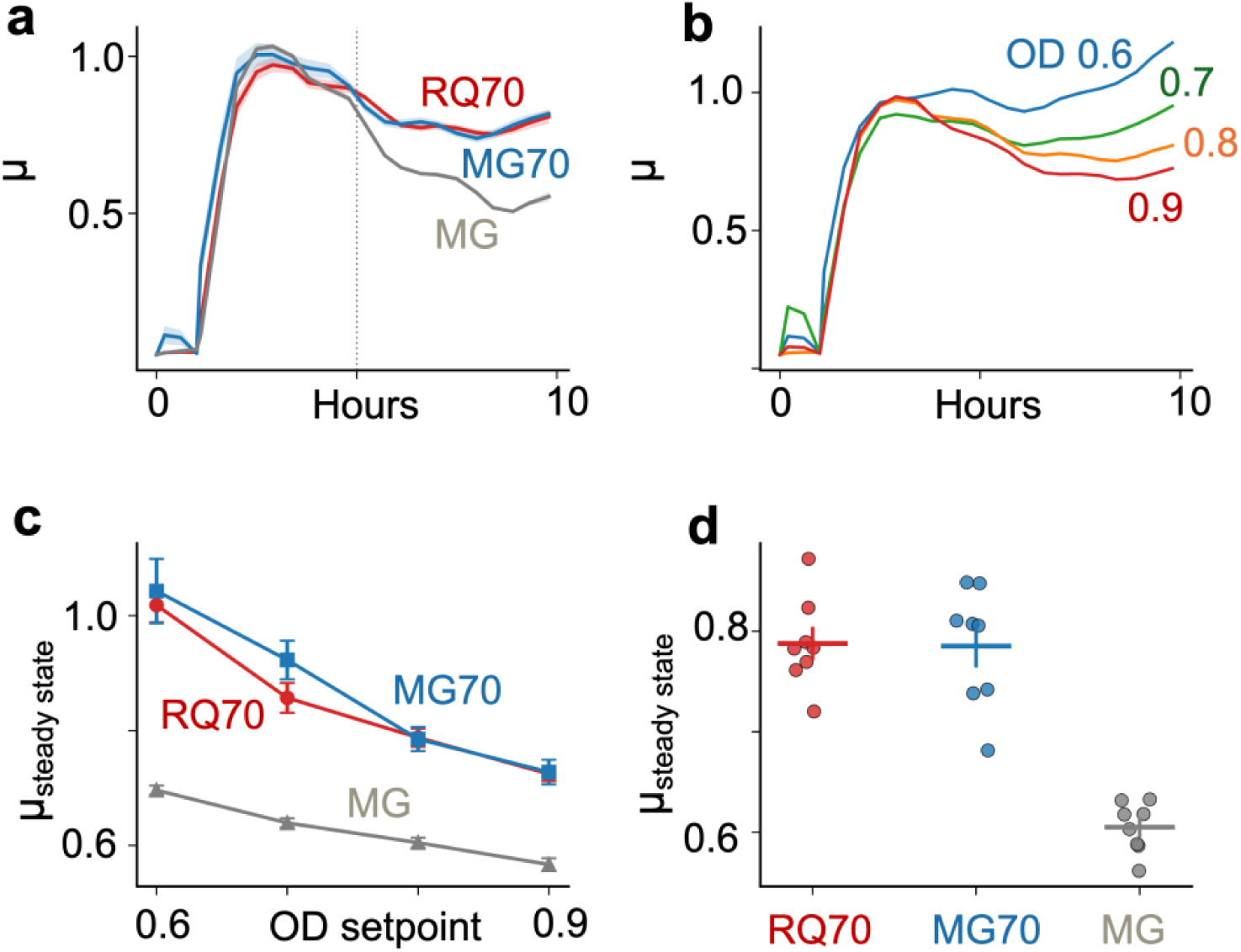
Turbidostat continuous culture reveals convergent steady-state growth rates between RQ70 and MG70 controls. **(a)** Growth rate time series at OD setpoint = 0.8 (mean ± s.e.m.; n = 8 biological replicates). Dotted line indicates the start of the steady-state window (t = 5 h). **(b)** RQ70 growth rate time series across OD setpoints 0.6–0.9. **(c)** Steady-state growth rate as a function of OD setpoint (mean ± s.e.m., t = 5–10 h). RQ70 and the MG70 control are indistinguishable across all setpoints, while both exceed the ancestor. **(d)** Individual well growth rates at OD setpoint = 0.8 (t = 5–10 h); horizontal bars indicate mean ± s.e.m. All cultures were grown in LB at 37 °C without antibiotics or inducers.

**Supplementary Figure 10.**
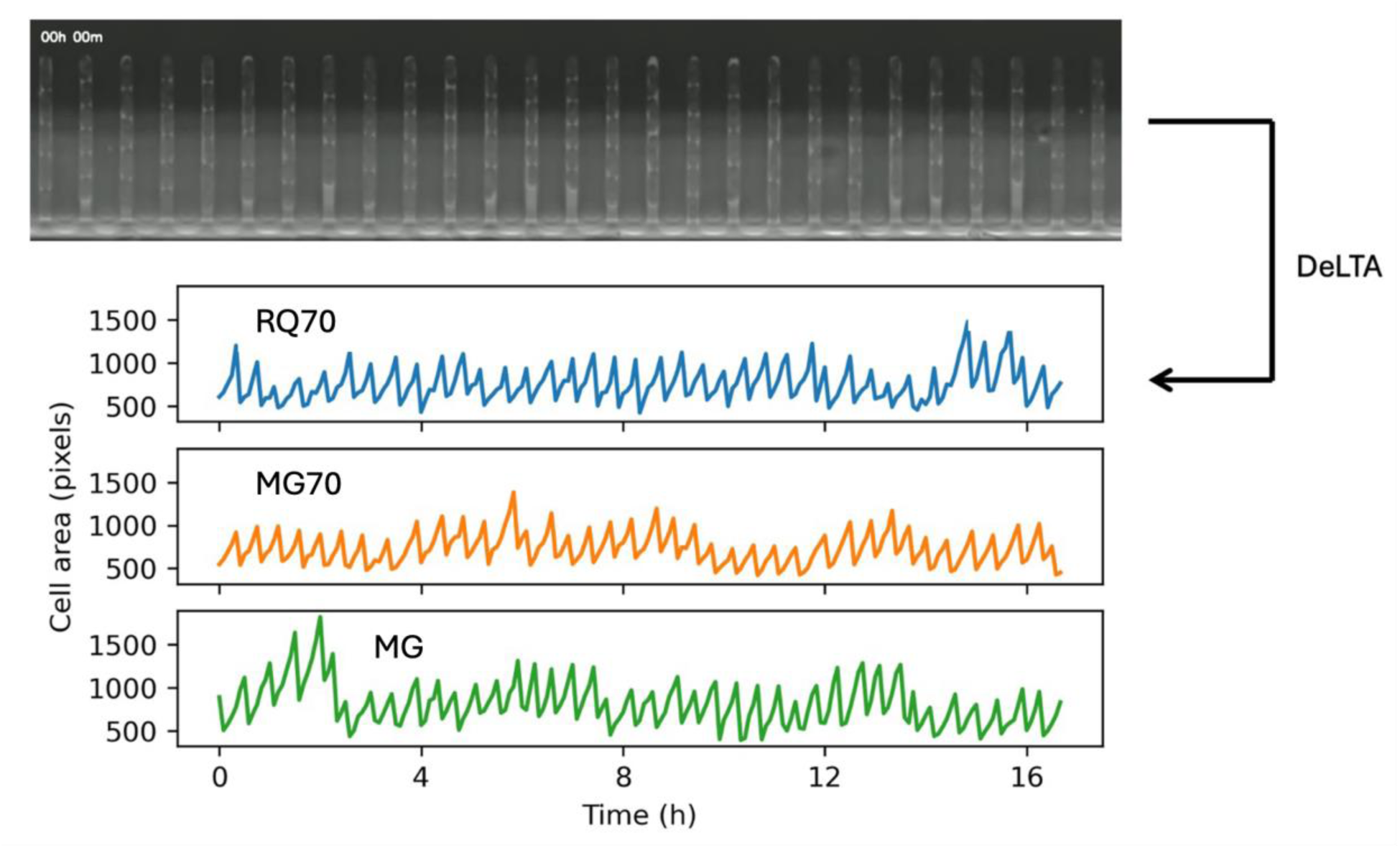
DeLTA segments cells and measures area over time in a mother machine. Representative phase-contrast field of the microfluidic mother machine at t = 0 (top), in which individual cells are confined in single-file growth channels under continuous medium replacement. Cell outlines were segmented and tracked using DeLTA, yielding single-cell cross-sectional area traces over time for RQ70 (blue), MG70 (orange), and the ancestral MG1655+FHr strain (green). The characteristic sawtooth pattern reflects successive rounds of elongation and division, from which single-cell elongation rates were derived (Supplementary Figure 11). Cells were grown in LB at 37 °C.

**Supplementary Figure 11.**
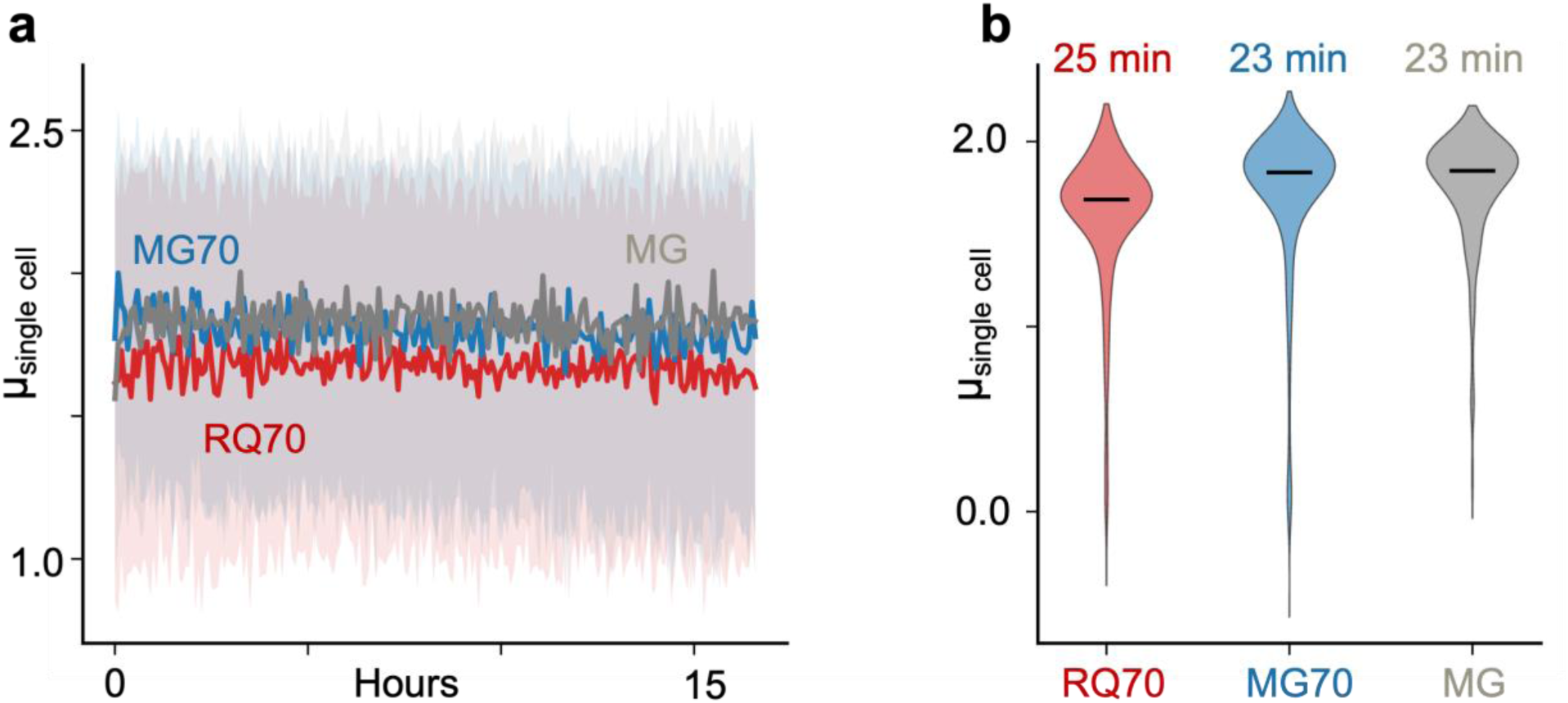
Growth rates over time of RQ70, MG70, and ancestral MG in a mother machine as calculated from cell area over time. Median growth rate time series **(a)** and distributions **(b)**. Numbers above distributions in **(b)** are doubling times calculated from median growth rate.

**Supplementary Figure 12.**
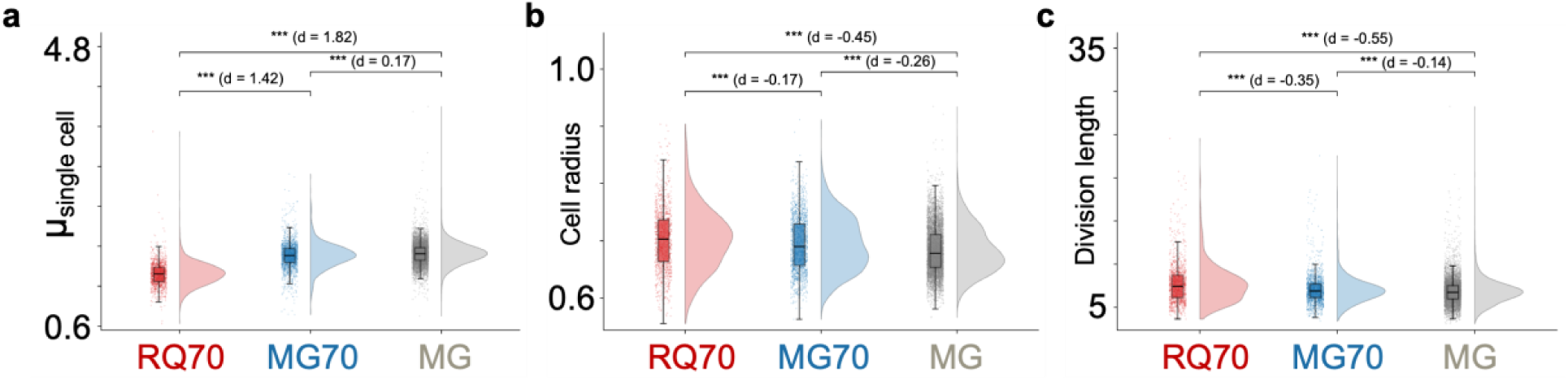
Single-cell morphology and elongation rate of RQ70 and MG70 on agar pads. Distributions of single-cell growth rate (hr^-1^) **(a)**, cell radius (µm) **(b)**, and division length threshold (µm) **(c)** for RQ70 (red, n = 1,183 cells), MG70 (blue, n = 1,683 cells), and the ancestral MG strain (gray, n = 6,223 cells) imaged on LB agar pads at 37 °C.

**Supplementary Figure 13.**
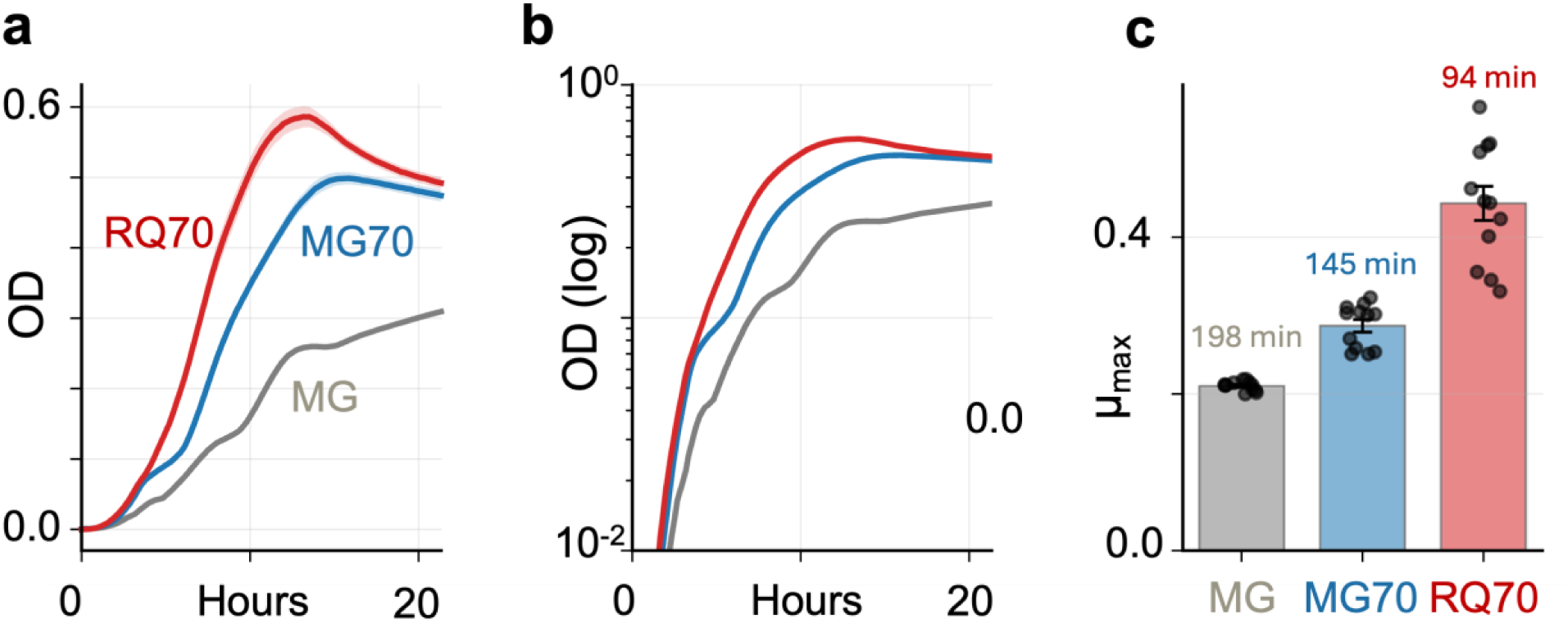
Growth kinetics of RQ70, MG70, and ancestral MG in M9 minimal medium supplemented with 0.4% glucose. Blank-subtracted OD600 growth curves on linear (A) and logarithmic (B) axes, and maximum specific growth rate from a logistic-with-lag fit (C), for RQ70 (red), MG70 (blue), and the ancestral MG1655+FHr strain (gray) in M9 + 0.4% glucose at 37 °C. Wells contained no plasmid, kanamycin, or inducer (n = 12 per strain). Absolute growth rates were substantially lower than in LB for all strains, but the rank order was preserved: RQ70 grew fastest, followed by MG70 and then the ancestor, with corresponding doubling times shown above each bar. RQ70 also established fastest and reached the highest final yield. Curves show mean ± s.e.m; bars show mean ± s.e.m. with individual replicates overlaid.

**Supplementary Figure 14.**
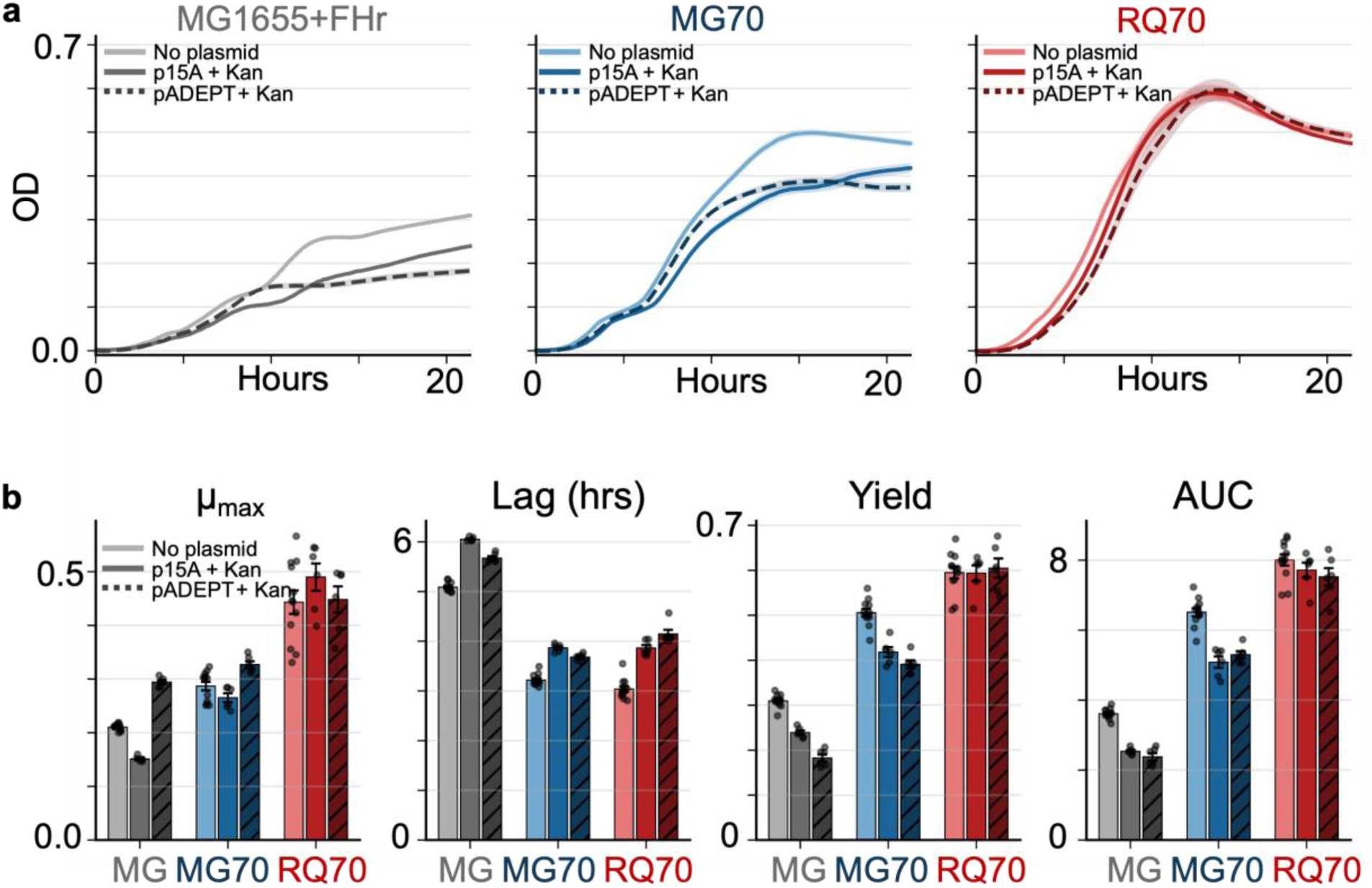
RQ70 shows smaller growth effects in the plasmid-kanamycin challenge than MG70 or the ancestor. (**a**) Blank-subtracted OD600 growth curves in M9 + 0.4% glucose for the ancestral MG1655+FHr (gray, left), MG70 (blue, middle), and RQ70 (red, right), each carrying no plasmid (M9 only, n = 12), the medium-copy p15A plasmid with kanamycin (n = 6), or the high-copy pADEPT plasmid with kanamycin (n = 6). (**b**) Growth rate from a logistic-with-lag fit, lag time (time to OD600 = 0.05), yield (maximum OD600), and integrated growth (AUC, OD*hr) for each strain–plasmid combination; shading denotes plasmid condition and hue denotes strain.

**Supplementary Figure 15.**
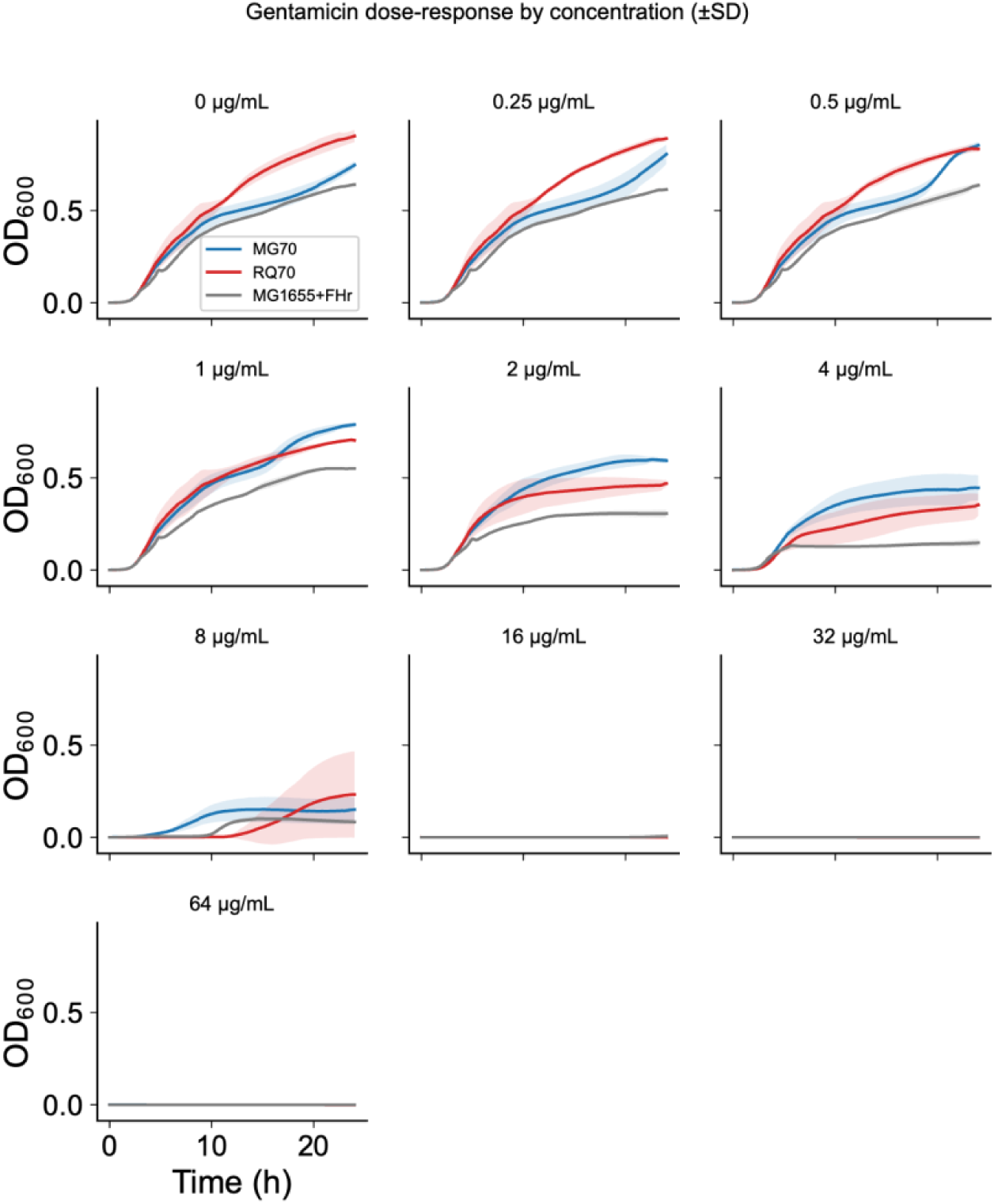
RQ70 shows delayed outgrowth at intermediate-to-high gentamicin concentrations. OD600 growth curves for RQ70 (red), MG70 (blue), and the ancestral MG1655+FHr strain (gray) in LB at 37 °C across a two-fold gentamicin dilution series (0–64 µg/mL), with one panel per concentration. No strain grew at or above 16 µg/mL, placing the MIC at 16 µg/mL for all three strains. At sub-MIC concentrations the strains separated: RQ70 grew fastest in the absence of drug, but at intermediate-to-high sub-MIC doses (2–8 µg/mL) it showed a prolonged lag followed by delayed exponential escape and was overtaken by MG70, which was the most drug-tolerant of the three.

**Supplementary Figure 16.**
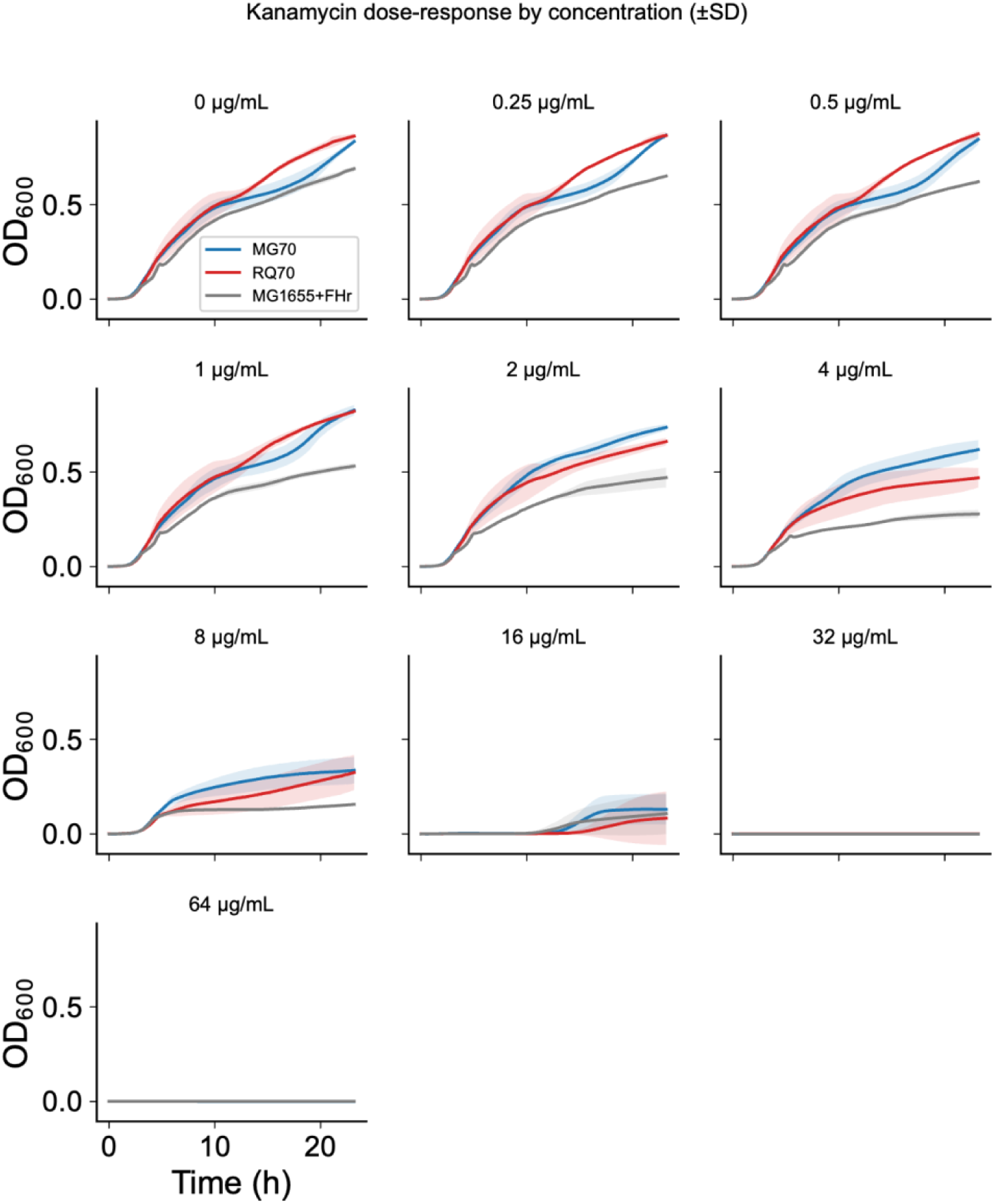
RQ70 shows delayed and reduced outgrowth at intermediate-to-high kanamycin concentrations. OD600 growth curves for RQ70 (red), MG70 (blue), and the ancestral MG1655+FHr strain (gray) in LB at 37 °C across a two-fold kanamycin dilution series (0–64 µg/mL), with one panel per concentration. Growth was detectable up to 16 µg/mL and fully suppressed at and above 32 µg/mL. As with gentamicin, the growth-rate ranking observed in drug-free medium inverted at intermediate-to-high sub-MIC concentrations (4–16 µg/mL), where MG70 sustained the greatest growth and RQ70 showed delayed and reduced outgrowth relative to the evolved control.

**Supplementary Figure 17.**
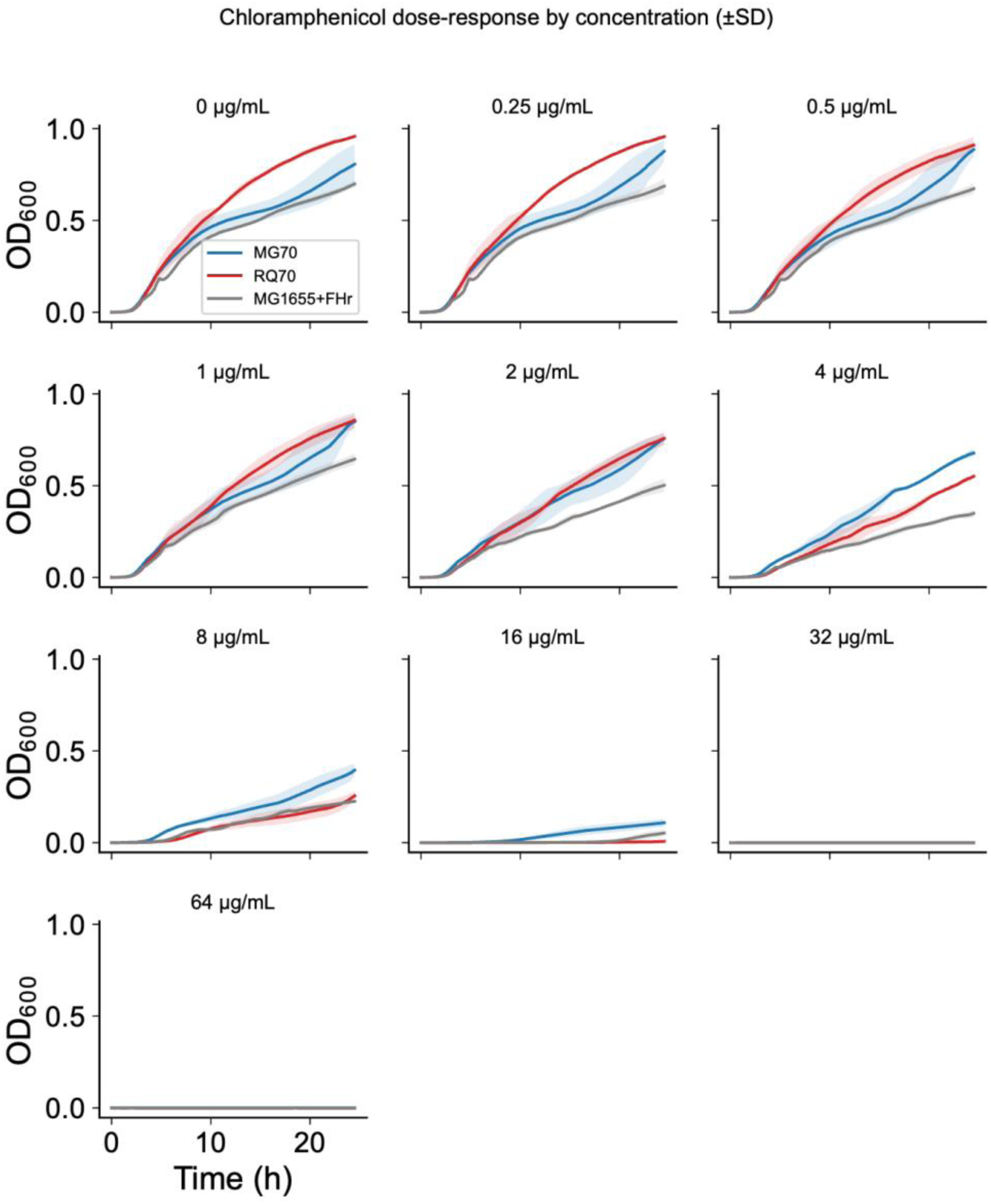
RQ70 shows delayed and reduced outgrowth at intermediate-to-high chloramphenicol concentrations. OD600 growth curves for RQ70 (red), MG70 (blue), and the ancestral MG1655+FHr strain (gray) in LB at 37 °C across a two-fold chloramphenicol dilution series (0–64 µg/mL), with one panel per concentration. Growth was fully suppressed at and above 32 µg/mL for all strains; at 16 µg/mL only MG70 showed appreciable outgrowth. RQ70 grew fastest in drug-free medium but lost its advantage from 4 µg/mL upward, falling to or below the ancestor at 8–16 µg/mL, while MG70 remained the most tolerant strain across the sub-MIC range.

**Supplementary Table 1.** Parameter values used in this study.

| Parameter | Description | Value |
| --- | --- | --- |
| $\mu_{max}$ | Maximum cell growth rate | 1.0 (fast), 0.5 (slow) |
| $\delta_{max}$ | Maximum RQ-mediated cell death rate | 1.0 |
| $K_1$ | pTarget copy-number rescue constant | 0.7 |
| $n$ | Hill coefficient for death rate | 3 |
| $\theta$ | pTarget replication rate | 1.0 |
| $K_2$ | pTarget replication Michaelis constant | 0.1 |
| $d_P$ | Cas9-mediate plasmid digestion rate | 1.0 |
| $P_{threshold}$ | pTarget replication threshold | 0.1 |
| $k_E$ | Maximum Cas9 expression rate | 0 (off), 0.1 (on) |
| $d_E$ | Cas9 degradation rate | 0.05 |

The model is non-dimensionalized: $\mu_{eff} = \mu_{max}(1 - N)$ means N is scaled to a carrying capacity of 1, so N, P, and E are all in arbitrary units and the rate constants are in $h^{-1}$ . The initial conditions for all circuit function simulations are: $Y_0 = [N, P, E] = [0.001, 1.0, 0.0]$ .

**Supplementary Table 2.** Per-figure *μ_max_* and *k_E_*settings.

| Figure | $\mu_{max}$ | $k_E$ | Integration |
| --- | --- | --- | --- |
| 1c | 1.0 (fast), 0.5 (slow) | 0.0 (off), 0.1 (on) | 0-16 hr |
| 1d | [0, 1.25] | 0.0, 0.1, 0.3, 0.6 | 0-16 hr |
| 2a-c | 1.0 | 0.0 (off), 0.1 (on) | 0-24 hr, 1:1,000 dilution, 24-40 hr |
| 2f | 1.0 | 0.0 (off), 0.1 (on) | 3x12 hr phases, 1:1,000 dilution between each |
| 2g | 1.0 | 0.0 (off), 0.1 (on) | 0-24 hr, 1:1,000 dilution, 24-36 hr |

**Supplementary Table 3.**
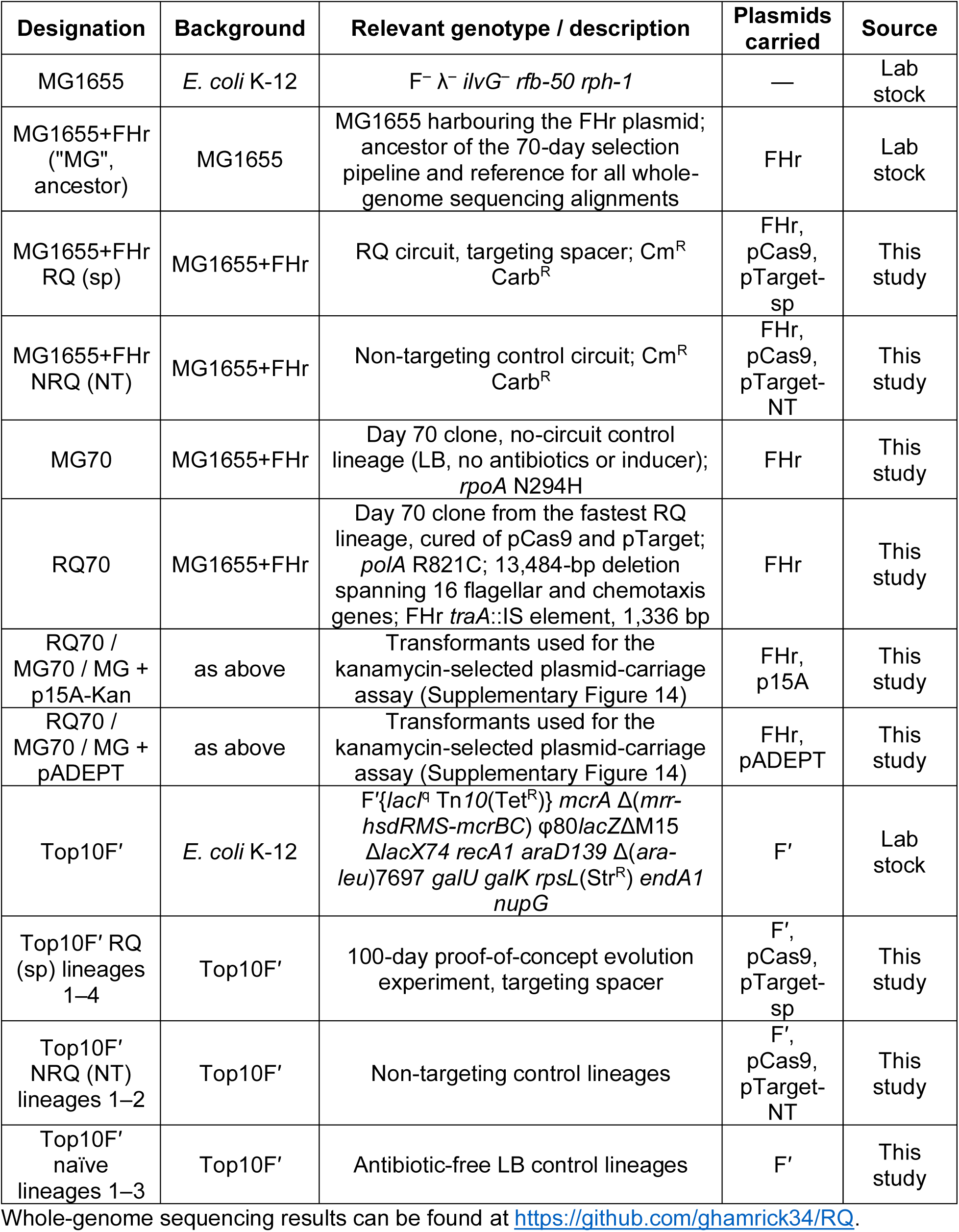
Bacterial strains used in this study.

**Supplementary Table 4.** Plasmids used in this study.

| Plasmid | Replication origin (copy number) | Selectable marker | Key features | Source |
| --- | --- | --- | --- | --- |
| pCas9 | SC101 (low) | *cat* (Cm<sup>R</sup>) | *Ptet* → *cas9* fused to an *ssrA* degradation tag; *tetR* | This study |
| pTarget-sp | pUC (high) | *bla* (Amp<sup>R</sup>/Carb<sup>R</sup>) | *Ptet* → guide RNA targeting a non-coding region on pTarget itself; constitutively expressed *bla* serves as the essential survival gene | This study |
| pTarget-NT | pUC (high) | *bla* | Mismatched, non-targeting guide RNA; otherwise isogenic to pTarget-sp | This study |
| pTarget-ColE1 | ColE1 (medium) | *bla* | Earlier circuit iteration; insufficient β-lactamase expression caused circuit loss (**Supplementary Figure 2A**). Not used beyond backbone optimization | This study |
| FHr | F / RepFIA (1–2) | Tn10 (Tet<sup>R</sup>) | Engineered derivative of the *E. coli* F fertility factor; with reduced mobilization efficiency by the F relaxase and a deletion of the *traS* gene involved in entry exclusion | 64 |
| F' (resident in Top10F') | F / RepFIA (1–2) | Tn10 (Tet<sup>R</sup>) | F'(*lacI*<sup>q</sup>); resident episome of Top10F' | Invitrogen |
| p15A | p15A (medium) | *aph* (Kan<sup>R</sup>) | Medium-copy comparator for the plasmid-burden assay | Lab stock |
| pADEPT | pUC (high) | *aph* (Kan<sup>R</sup>) | High-copy comparator for the plasmid-burden assay | 42 |

Plasmid maps for pCas9, pTarget-sp, pTarget-NT, pTarget-ColE1, p15A, and pADEPT can be found in the supplement at https://github.com/ghamrick34/RQ.

